# Integrating NMR restraints into coarse-grained simulations: toward accurate conformational ensembles of complex protein systems

**DOI:** 10.64898/2025.12.22.695971

**Authors:** Mina Cullen, Carmen Biancaniello, Katerina Taškova, Vedran Miletić, Davide Mercadante, Alfonso De Simone

## Abstract

Structural dynamics play critical roles for the biological activity of protein molecules. Characterising the inherent conformational landscapes of these macromolecules remains a major experimental and computational challenge, particularly for heterogeneous and transient systems such as intrinsically disordered proteins, membrane-associated assemblies and disordered fuzzy coats of amyloid aggregates. In this context, coarse-grained (CG) molecular dynamics simulations have enabled accessing to extended timescales and large system sizes, however, their reduced resolution and simplified interaction potentials often limit the structural accuracy.

Here, we introduce Martini3-NMR, an integrative framework that incorporates nuclear magnetic resonance (NMR) observables directly into CG protein force fields. Using artificial neural networks to model NMR chemical shifts at the CG level, and integrating these data with NOE restraints, we define an approach to significantly enhance the accuracy of CG simulations while maintaining their elevated sampling efficiency, thereby resulting in a substantially improved description of protein conformational ensembles. We demonstrate the broad applicability of Martini3-NMR by generating CG ensembles for a diverse range of systems and complex processes such as protein folding from misfolded states, oligomer disassembly within lipid bilayers and the topology of disordered fuzzy regions at amyloid fibril surfaces, which were found to display condensate-like structural and dynamical properties.

By enabling an experimentally driven and computationally efficient exploration of protein conformational landscapes, Martini3-NMR provides a novel general framework for investigating dynamic, heterogeneous and multiscale biomolecular processes. This approach opens to significant new opportunities for extending CG simulations towards a more quantitative understanding of the relationship between molecular structure, dynamics and biological function.

## Introduction

Structural fluctuations are fundamental determinants of the biological activity of proteins, as these biomolecules populate broad ensembles of interconverting conformations, encompassing both folded and intrinsically disordered native states.^1^ These structural dynamics span a wide range of timescales, from picoseconds to seconds, and enable proteins to access the specific conformations required for diverse biological processes, including enzymatic catalysis,^2^ oligomeric assembly,^3^ molecular recognition and signalling,^4^ and others. A detailed characterisation of these motions is critical for elucidating how proteins engage their interaction partners and dynamically access distinct functional states. Characterising conformationally heterogeneous proteins remains a major experimental challenge, particularly when the relevant states are sparsely populated or transient in nature. This major limitation hampers the study of biochemical processes that are governed by low-population protein states accessed only transiently through their inherent conformational energy landscapes. Such intermediates often play a decisive role in processes such as protein folding, conformational switching, allosteric regulation, enzymatic turnover and the assembly or disassembly of biomolecular complexes, however, their short lifetimes, intrinsic dynamics and minimal abundance render them exceptionally challenging to characterise at high resolution using conventional structural biology approaches.

When studying intermediate states that exist in dynamic equilibrium with ground-state conformations, nuclear magnetic resonance (NMR) spectroscopy has emerged as a uniquely powerful experimental approach, enabling access to a broad range of timescales and spatial resolutions.^5,6^ This technique can provide detailed structural, thermodynamic, and kinetic information through the detection of time- and ensemble-averaged observables that are inherently sensitive to conformational fluctuations. Through its evolution into solution-state and solid-state methodologies, NMR has progressively expanded its scope to encompass a wide range of systems, including cytosolic proteins, membrane proteins,^7,8^ insoluble amyloid fibrils,^9–12^ megadalton complexes^13,14^ and biomolecular condensates.^15,16^ In parallel, theoretical approaches have become increasingly important for the investigation of protein structural fluctuations at high resolution, with molecular dynamics (MD) simulations assuming a central role. When integrated with NMR data as experimental restraints, the intrinsic limitations of MD force fields can be substantially mitigated, resulting in improved accuracy in the determination of dynamic conformational ensembles.^5,17–19^ Over the past decades, several powerful refinement strategies have been proposed to incorporate NMR restraints into full-atom MD frameworks, achieving remarkable success in improving structural accuracy and capturing local and global dynamical properties.^19–23^ Nevertheless, the inherent limitations of all-atom MD in sampling large conformational phase spaces on biologically relevant timescales restrict its applicability to the study of highly heterogeneous and dynamic systems.

Here, we present a study reporting substantial advance in the conformational sampling of proteins by introducing NMR restraints into coarse-grained (CG) models within a simulation framework designated as *Martini3-NMR*. CG simulations offer strategic advantage for exploring extensive conformational landscapes and accessing long timescales that are typically inaccessible to all-atom approaches. By integrating the experimental accuracy of NMR restraints with the enhanced sampling efficiency afforded by CG simulations, we show that complex biochemical processes can be characterised with unprecedented level of detail, thereby extending the scope and applicability of both techniques. We provide additional support for these conclusions by showing that NMR chemical shifts (CS) and Nuclear Overhauser effect (NOE) restraints applied within the Martini3 CG model enable accurate sampling of the conformational space of both soluble and membrane proteins, while also capturing complex processes such as the functional disassembly of oligomeric proteins within the lipid bilayers and the delineation of the topological and structural properties of disordered fuzzy coats in amyloid fibrils.

## Results

### NMR chemical shift calculations in coarse-grained protein structures

We developed artificial neural networks (ANNs) to predict NMR CS of proteins from CG structural models. The ultimate goal of our work is to incorporate this model, namely NapShift-CG, as an experimental NMR-derived restraining potential into CG simulations, thereby enhancing both sampling efficiency and the structural accuracy of the resulting conformational ensembles. ANNs were generated to account for the topology of the Martini3 CG force field by deriving internal parameters from the Cartesian coordinates of the backbone (BB) and sidechain (SC) beads (Figure 1a). More specifically, for each residue of the protein, NapShift-CG employs three bond angles centred on a BB (α, β, γ) and two dihedral angles involving the flanking BBs (θ_1_ and θ_2_). This topological definition makes NapShift-CG applicable to other Martini force fields including dry Martini and pol-Martini, upon re-parametrization of the ANN. The method was designed to account for NMR CS of six distinct protein atoms from the main chain (N, C_α_, C, HN and H_α_) and side chains (C_β_). Instead of generating six individual ANNs where each protein atom is treated separately, as previously developed for the full-atom NapShift predictor,^24^ in NapShift-CG all six atom types are trained simultaneously. We evaluated different structural parameters derived from mono-, tri-, penta-, and heptapeptides to contribute input vectors for ANN training (Figure 1), and found that tripeptides generate the best compromise between accuracy of NapShift-CG and computational complexity. ANNs training was performed using 2,986 protein structures for which CS assignments were available. A set of 250 entries from the initial database was not included in the training dataset and was used as a validation dataset (see the Supplemental Data spreadsheet).^24^

**Figure 1.**
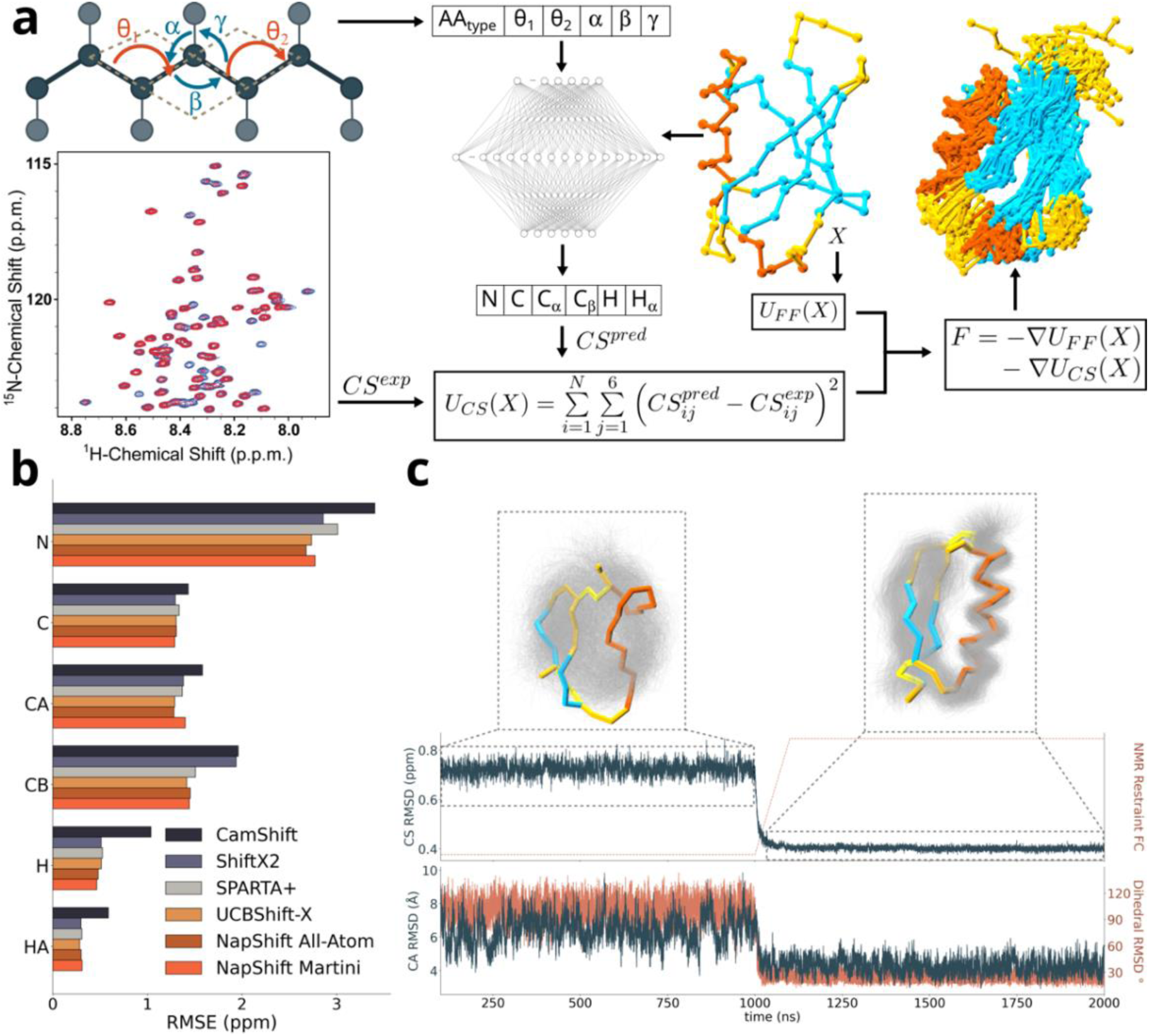
Conceptualisation, implementation and testing of Martini3-NMR. (a) Coarse-grained heptapeptide mapped with Martini3. Backbone and sidechain beads are in dark and light grey, respectively. The dihedrals θ_1_ and θ_2_ and the angles α, β, and γ are the Napshift-CG inputs. Napshift-CG predicts chemical shifts (CS) for backbone protein atoms N, C, C_α_, C_β_, H and H_α_. Predicted (CS^pred^) and experimental (CS^exp^) chemical shifts are compared at each simulation step, and their difference is used to correct the discrepancy by adjusting the forces acting on the beads and to obtain CG protein ensembles, in line with experimental NMR data. On the right-hand side the CG experimental structure and the obtained ensemble of ubiquitin are coloured by secondary structure (α-helices – orange, β-strands – cyan, loops – yellow). The system was simulated with K_CS_ = 0 for 1 μs, after which K_CS_ was increased to 25 over 100ns, then simulated for a further 1 μs. The first 100 ns were discarded as equilibration time. (b) Root mean square error (RMSE) reporting on the predictive ability of each backbone atom by Napshift-CG and a series of all-atom predictors. (c) Simulation of a de novo designed mini-protein (PDB ID: 2ND3) before and after the application of NMR CS restraints. Top and bottom panel show the root mean square deviation (RMSD) in CS (top), dihedral and coordinates (bottom) space. The insets show the obtained ensembles (gray) before (left) and after (right) the application of the restraints. The ensemble centroids are overlaid on the ensemble and coloured by secondary structure as in (a).

With this architecture, the accuracy of NapShift-CG resulted of similar magnitude to that of methods based on full-atom structures (Figure 1b). This result holds strategic relevance, as it established that CS of proteins can be predicted from CG models with state-of-the-art accuracy, thereby opening to the way to several advanced applications in structural refinement and MD simulations.

### Definition of NMR-restrained coarse-grained simulations of proteins

The architecture of NapShift-CG was based on structural parameters that are differentiable in the Cartesian space, thereby enabling the calculation of gradients and forces driving the simulations toward an improved match between computed and experimental CS (Figure 1b and Figure S1). CS restraints were applied to the Martini3 force field using the OpenMM software^25^ (see Methods), and were initially tested on a de novo designed mini-protein for which the experimental structure (PDB code: 2ND3) and CS (BMRB code: 26046) are available.^26^ When simulating this system, unrestrained Martini3 MD simulations provided a poor representation of its conformational sampling, reaching a C_α_ mean RMSD of 6.44 ± 0.045 Å and mean dihedral RMSD of 103.16° ± 0.62° from the experimental structure (Figure 1c). The introduction of CS restraints via a gradual increase of the applied force was found to induce a rapid reduction in the CS RMSD, indicating that the implementation of the NapShift-CG derivatives was effective. Notably, the reduction in CS RMSD was found to correspond to an improvement of the conformations sampled during the MD simulations, with mean values of C_α_ and dihedral RMSD plateauing at 4.22 ± 0.02 Å and 24.18° ± 0.21°, respectively. This result is of fundamental significance, as it demonstrates that the incorporation of NMR restraints into CG force fields, such as Martini3, can dramatically enhance the ensemble quality of simplified models, particularly with respect to the dihedral angles of the protein backbone.

We next evaluated NapShift-CG on ubiquitin, a protein system featuring a more complex topology that includes a single β-sheet of five strands (in mixed parallel and antiparallel topology), an α-helix (residues 23-34) and a 3₁₀-helix (residues 56-59).^27^ Using unrestrained Martini3, CG MD simulations are unable to retain the Ub structure, with mean C_α_ and dihedral RMSD of 13.47 ± 0.12 Å and 104.49° ± 0.40°, respectively (Figure S2). When applying NapShift-CG NMR restraints, the conformational space explored improved considerably with respect to the dihedral angles (with mean RMSD of 47.37° ± 0.19°), however, the simulations could not accurately capture tertiary contacts between regions that are distant in the protein sequence, leading to a mean C_α_ RMSD of 14.02 ± 0.13 Å (Figure 2a). We therefore postulated that CS restraints in CG simulations should be combined with additional NMR observables that encode the topological features of protein structure (Figure S2). To this end, we selected distance information derived from NMR NOEs as the optimal complement to CS (see Methods). The combination of CS and NOE restraints, here after described as Martini3-NMR, resulted in a significant improvement of the quality of the Ub structures sampled by CG simulations with mean values of C_α_ and dihedral RMSD of 3.20° ± 0.01° Å and 44.88° ± 0.12°. Using only NOEs instead led to structural ensembles sampled with Martini3 that were of poor quality, particularly with respect to dihedral angles (Figure S2b-c).

**Figure 2.**
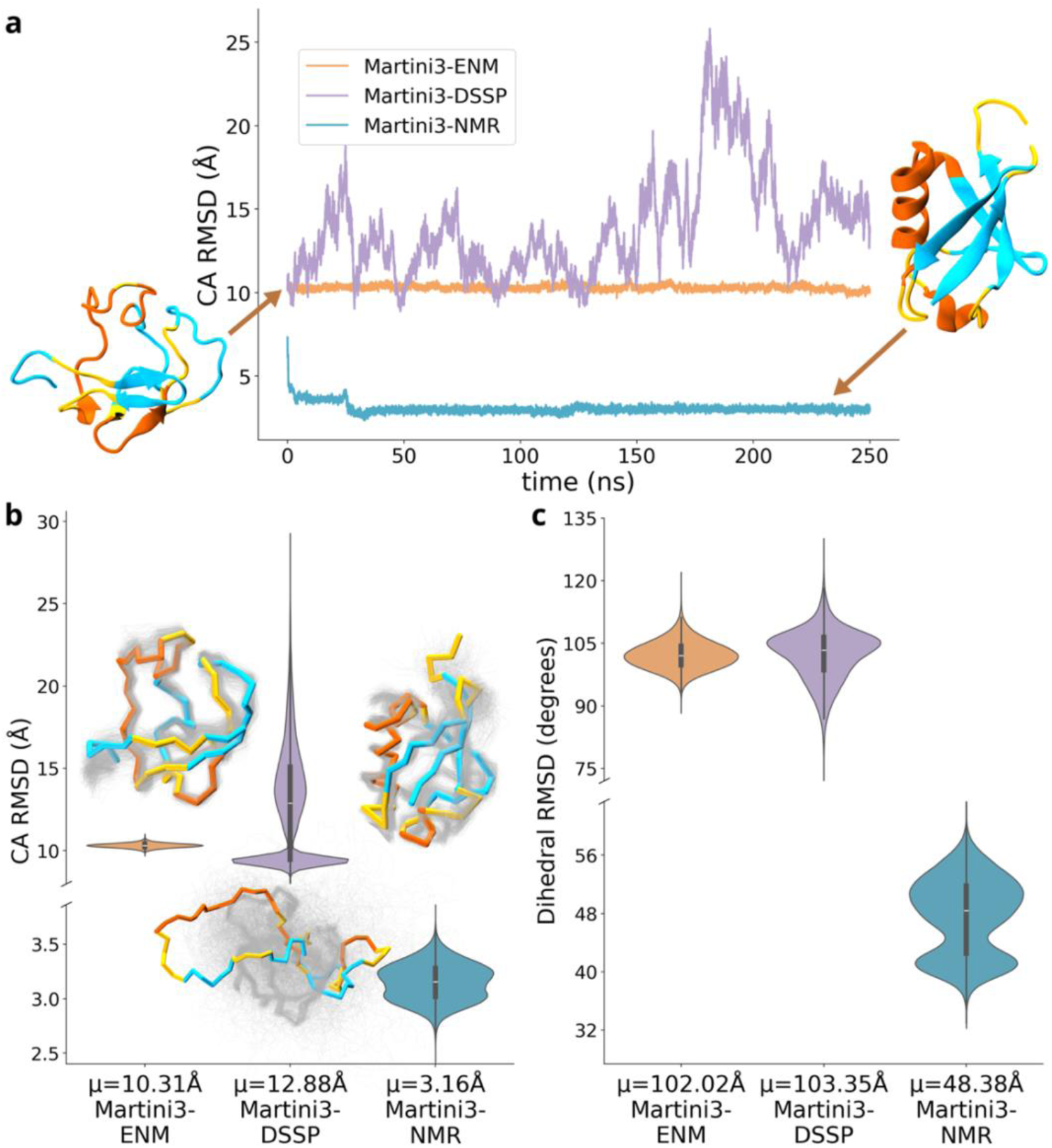
Effects of CS restraints on a partially unfolded protein. (a) Time trace of the C_α_ root mean square deviation obtained from simulations of ubiquitin with the initial conformation partially unfolded (C_α_ RMSD = 11.83 Å). (b-c) Coordinates (b) and dihedral (c) RMSD distributions for the simulations shown in (a). The obtained ensembles are shown above each violin plot with the distribution means (μ) reported in the tick labels.

The key aspect of Martini3-NMR lies in its ability to tailor CG-MD to specific biochemical processes, as the incorporation of system-specific NMR restraints enables the generation of ensembles that are also sensitive to specific experimental conditions under which the NMR data are acquired. Another distinct advantage of the proposed method is that it overcomes the limitations of CG approaches that restrict the conformational space of a protein to its initial coordinates, such as for example elastic network models (ENM) or Martini3-DSSP.^28^ We illustrate this fundamental point by performing CG simulations initiated from a misfolded Ub structure (Figure 2), exhibiting a C_α_ RMSD of 11.83 Å relative to the native state. When running CG simulations with elastic networks (Martini3-ENM), the conformations remained essentially frozen to the initial distorted state (C_α_ RMSD of 10.31 ± 0.01 Å), whereas the employment of Martini3-DSSP generated a variety of misfolded conformations with values of C_α_ RMSD varying from 7.99 Å to 28.79 Å along the trajectory. By contrast, when using Martini3-NMR, the system rapidly folded into the native structure, plateauing at a mean C_α_ RMSD of 3.16 ± 0.01 Å. This result highlights the fundamental improvement introduced by NMR-restrained CG simulations, which do not simply constrain the conformational space of the CG protein model but rather enhance the force field accuracy by steering the conformational ensemble toward states that are more consistent with experimental data.

Having established the optimal integration of sparse NMR data to restrain CG force fields, we tested Martini3-NMR on a variety of globular proteins of increasing structural complexity (Figure 3a). First, we tested this method on KRAS, a protein that, similarly to Ub, shows a mixed α-helical/β-sheet topology, but significantly larger than ubiquitin. KRAS is a GTPase that regulates cellular pathways involved in signalling for growth, proliferation, and differentiation. It is also the most frequently mutated oncogene in human cancer, making it an important system for structural biology. Martini3-NMR was able to retain the overall fold of KRAS with average C_α_ and dihedral RMSD values of 4.56 ± 0.03 Å and 41.53° ± 0.20°, respectively, with significant improvement compared to with Martini3 and Martini3-DSSP (Figures 3a, S3 and S4). A similar high-quality conformational ensemble was generated for the SH3 tandem of the human KIN protein (C_α_ RMSD of 2.07 ± 0.01 Å and mean dihedral RMSD of 38.95° ± 0.33°, Figures 3a, S3 and S4), a system of increased structural complexity owing to the presence of two intertwined domains. We also generated the NMR-guided CG ensemble of the human leptin, a multi-potency cytokine that regulates various physiological functions and whose structure is composed of a mainly five-helix bundle structure with two hydrophobic cores. The method generated an exceptionally good ensemble of leptin with mean C_α_ RMSD of 3.38 ± 0.02 Å and mean dihedral RMSD of 32.91° ± 0.23° (Figures 3a, S3 and S4). The ensemble was also able to capture the conformational properties of intrinsically disordered regions (IDRs) in the loop connecting helices A and B as well as the flexible helix E residing in the long loop connecting helices C and D, by improving the conformational space with respect to the unrestrained simulations (Figure S5). Both dynamic regions are essential for the conformational recognition of leptin by its receptor, and their properties were accurately captured by Martini3-NMR, suggesting that the mechanisms underlying key signalling pathways can be investigated using this hybrid CG-NMR force field.

**Figure 3.**
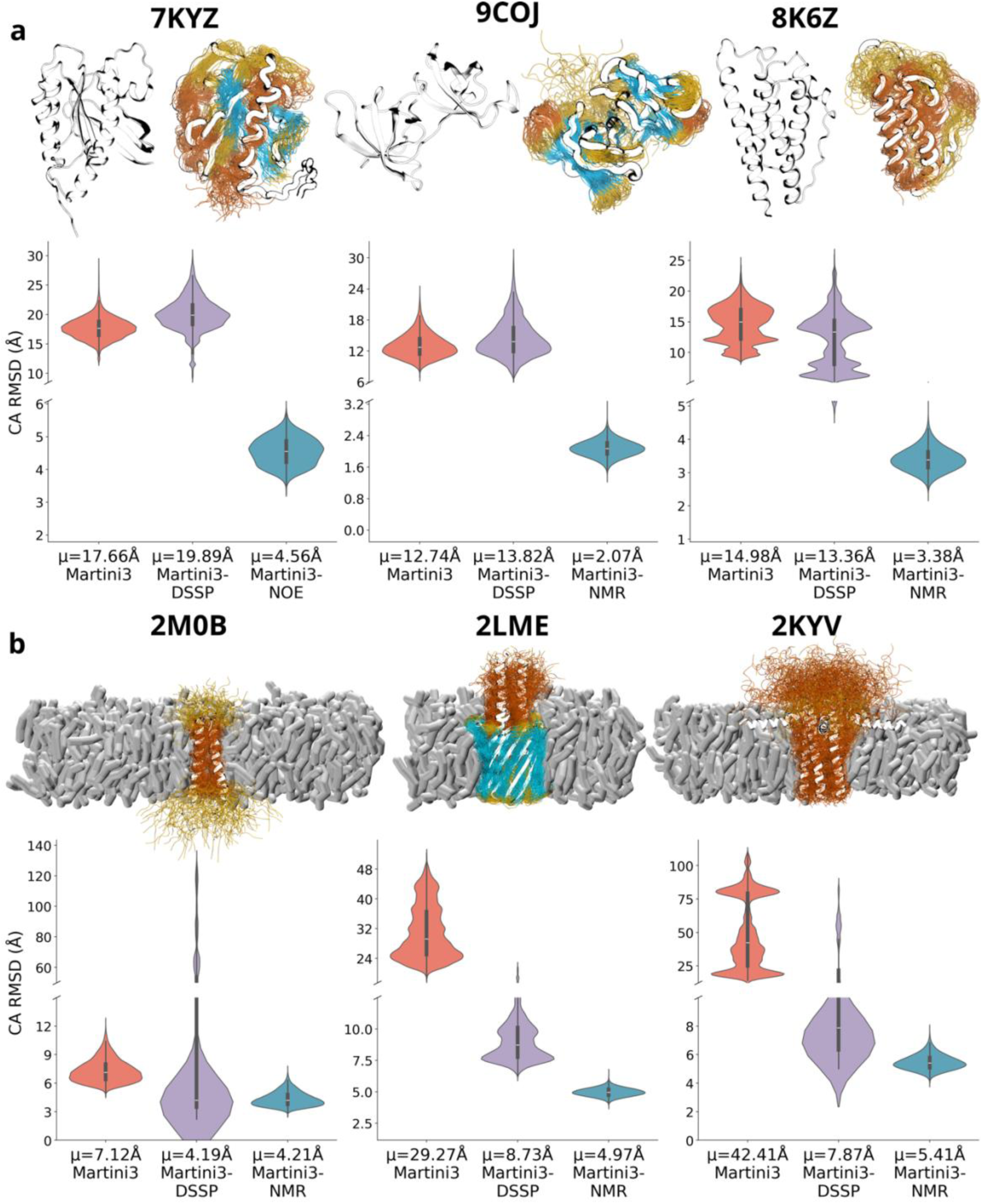
Benchmarking Martini3-NMR on soluble and membrane proteins. (a-b) Representative ensembles and root mean square deviation (RMSD) distributions in the Cartesian space (C_α_) of soluble (a) and membrane (b) proteins. (a) Soluble proteins from left to right are KRAS, the SH3 tandem domains of the human KIN protein and a human leptin. α-helices, β-strands and loops are shown in orange, cyan and yellow, respectively. On the left panel the experimental structures are reported, while on the right panel the ensembles overlaid on the centroid conformations are shown. (b) Membrane proteins from left to right are the single-span transmembrane helical domains of the human tyrosine kinase ErbB1, the transmembrane anchor domain of the bacterial autotransporter YadA, and the phospholamban pentamer. PDB codes are reported on top of the ensembles. The colour code for secondary structure is the same as in panel (a).

### Conformational ensembles of membrane proteins with Martini3-NMR

In addition to soluble proteins, we leveraged the capability of Martini3-NMR to model biological membranes and applied this approach on some relevant membrane-protein systems with increasing topological complexity (Figure 3b). First, we simulated the homodimeric transmembrane (TM) domain of the EGFR HER1, whose structure was originally resolved by solution NMR in micelles.^29^ The conformational transition between inactive and active dimer configurations of the HER1 TM domain controls receptor signalling in functional and pathological conditions such as cancer.^30^ We first modelled the structure of the TM domain by equilibrating it in a physiological lipid bilayer composed of 400 DLPC lipids (20 x 20 nm along the X and Y dimensions) as described in Thomasen *et al.*^31^ and to match experimental conditions.^32^ The simulations with Martini3-NMR sampled the dynamics of the inactive conformation of the TM dimeric domain by yielding a mean C_α_-RMSD of 4.21 ± 0.04 Å and mean dihedral RMSD of 43.07° ± 0.3° (Figure S4). The analysis of the inter-helical tilt angle revealed that the relative orientation of the two helices yields a dimer stably sampling the inactive state, with the most populated conformation matching the experimental value of the tilt angle. Nonetheless, the distribution of tilt angles shows that Martini3-NMR also samples a minor population at negative angle values, suggestive of a pathway leading to an active state (Figure S6).

We next investigated the TM domain of the Yersinia enterocolitica adhesin A (YadA), whose structure has been resolved in a lipid bilayer by solid-state NMR.^33^ This membrane domain exhibits a more complex topology, consisting of a homotrimer forming a 12-stranded membrane β-barrel, with each monomer contributing four antiparallel strands that assemble into a single symmetric pore. In this structure, each monomer also features an additional short N-terminal α-helix that extends into the aqueous phase (Figures 3b ,S3 and S4). CG simulations of YadA yielded a mean RMSD of 4.97 ± 0.02 Å for the β-barrel and 6.44 ± 0.03 Å when including the incomplete helical region. Although these RMSD values are higher than those obtained for other systems examined, they remain substantially lower than those produced using Martini3 and Martini3-DSSP, indicating a significant improvement introduced by the incorporation of NMR restraints (Figure 3b).

As an additional membrane protein system, we studied the pentameric assembly of the phospholamban protein (PLN) (Figure 3b and 4).^34^ This single-pass TM protein plays a critical physiological role by regulating the SERCA (sarcoplasmic/endoplasmic-reticulum Ca²⁺-ATPase) pump, thereby regulating cardiac muscle relaxation and contractility. In its physiological form, PLN exists in equilibrium between pentameric and monomeric states, with the oligomer believed to act as a reservoir for the inhibitory monomeric state, which is the form that directly interacts with SERCA.^35^ We generated the structural ensemble of the pentameric PLN using Martini3-NMR starting from a hybrid solution and solid-state NMR structure resolved in lipid bilayer.^33^ The ensemble exhibited high accuracy in modelling the TM region of the oligomer, as it successfully reproduced the orientations of the homopentameric helical bundle (residues 24 to 52), in close agreement with the full-atom ensemble previously obtained from orientational restraints and conveying the packing of the bundle identified as stabilising components of PLN multimeric state and relevant for the general design of stably packed TM helical domains (Figure 4a),^34,36^ ^37^ In contrast, the N-terminal membrane-surface helices resulted poorly associated with the surface of the lipid bilayer and as a consequence adopted random orientations. This behaviour is likely attributable to intrinsic limitations of the CG force field in modelling protein-membrane interactions of amphipathic helices placed on top of the lipid bilayer. This limitation could be mitigated through the inclusion of orientational restraints, which are currently not implemented in Martini3-NMR.^38 37^

**Figure 4.**
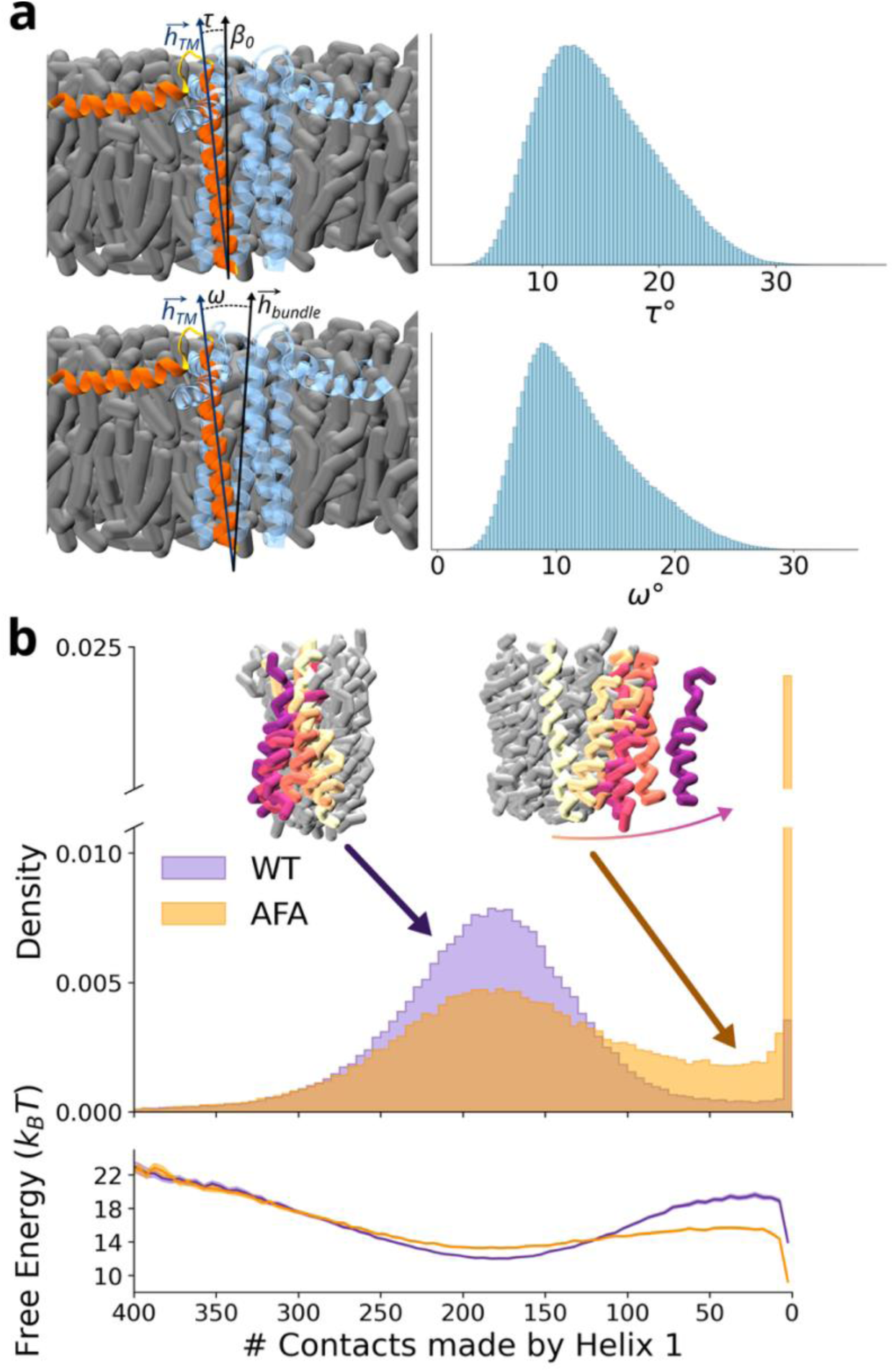
Martini3-NMR recapitulates the conformational features of the phospholamban pentameric state. (a) Phospholamban pentamer in a lipid membrane (gray). One of the helices, designated as helix 1, is shown in orange while the others are shown in blue, to highlight the angles τ and ω (top and bottom panels, respectively) used to describe the conformational dynamics of the pentamer as in Sanz-Hernandez et al.^38^ The distributions of τ and ω angles yielded by Martini3-NMR simulations are shown. These angles describe the orientation of phospholamban helices with respect to either the membrane norm (τ) or the helix bundle (ω). (b) To compute the distributions of contacts of helix 1 with the reminder of the pentamer, 200 simulations each of 500 ns of WT and AFA-mutant Phospholamban were performed, restraining the backbone bead of the last residue to an upper limit of 3.8 nm along the z-dimension. In the top panel, the number of contacts between helix 1 and the reminder of the pentamer molecules is shown in indigo orange for PLN^WT^ and PLN^AFA^, respectively. In the bottom panel the free energy landscape along the same reaction coordinate has been computed using the Boltzman equation.

In addition to describing the topological-dynamical properties of the pentamer barrel, the NMR-guided CG simulations enabled sampling of the process of monomer detachment from the pentamer assembly (Figure 4b). Several lines of evidence exist about the intrinsic stability of the pentameric state of PLN with respect to the isolated monomers.^34,39^ By mutating the three Cys residues of the TM region into Ala 36, Phe 41 and Ala 46 (PLN^AFA^),^40^ it is possible to invert the energy landscape and push the equilibrium toward the monomeric state. We therefore exploited the sampling ability of CG simulations to study the monomer-pentamer equilibrium in PLN^WT^ and PLN^AFA^ (see methods). The data showed a marked difference between the two sequences when sampling a reaction coordinate of monomer detachment. In particular, while the WT stably sampled the pentameric state, with minor events of detachment, PLN^AFA^ sampled numerous trajectories of monomer detachment from the four reminder molecules, experiencing 70% detachment events vs 15% of the WT. The analyses indicate that, along a reaction coordinate based on the number of contacts between the detaching monomer and the remainder of the molecules composing the pentamer, the free energy of PLN^WT^ bound state is lower than that of the detached state. The scenario is completely inverted for PLN^AFA^ where the free energy of a configuration featuring a detached monomer from the reminder four monomers is lower than that of the assembled pentamer (Figure 4b). In addition, the AFA triple mutation also reduces the energy barrier of detachment to 2.44 ± 0.19 kJ/mol from 7.63 ± 0.19 kJ/mol of the WT. Collectively these data show that Martini3-NMR is particularly suitable for studying relevant biochemical processes in membrane proteins, such as the disassembly of the PLN pentamer along the regulation process of SERCA.

### Elucidating the properties of amyloid fibrils and their fuzzy coats using NMR restrained CG

Given its suitability to investigate large molecular systems, we next assessed the ability of Martini3-NMR to simulate large protein assemblies such as amyloids. To this end, we simulated the protease-resistant fragment (residues 297-391) from the Alzheimer’s disease Tau core, assembled into a non-twisted amyloid (Figure 5a).^41^ ssNMR studies identified a rigid amyloid core composed of residues 305–357, with residues outside this region remaining highly flexible. Because the Tau_297–391_ fibrils showed no detectable twist in TEM, precluding the use of helical reconstruction methods, solid-state NMR was crucial for resolving their molecular structure. Our Martini3-NMR CG simulations retained the structural properties of the fibrillar region of the core (residues 305-357) with mean C_α_ RMSD of 5.64 ± 0.01 Å and dihedral RMSD of 57.47° ± 0.09° (Figure S7). When simulating with Martini3 or Martini3-DSSP the fibril core rapidly disassembled yielding non-viable simulations. In addition, beyond maintaining fibril stability, Martini3-NMR simulations were able to retain the network of key inter-sheet interactions, including Y310–V337, H329–D348, L325–V350, and L325–I354, which collectively lock the opposing β-sheets into a compact and highly stable amyloid core. The symmetric E338–E338 interface was however not well accounted, likely because protonation states of Glu residues were not defined in the featurisation of the ANN due to the lack of a sufficient amount of specific data (Figure 5b).

**Figure 5.**
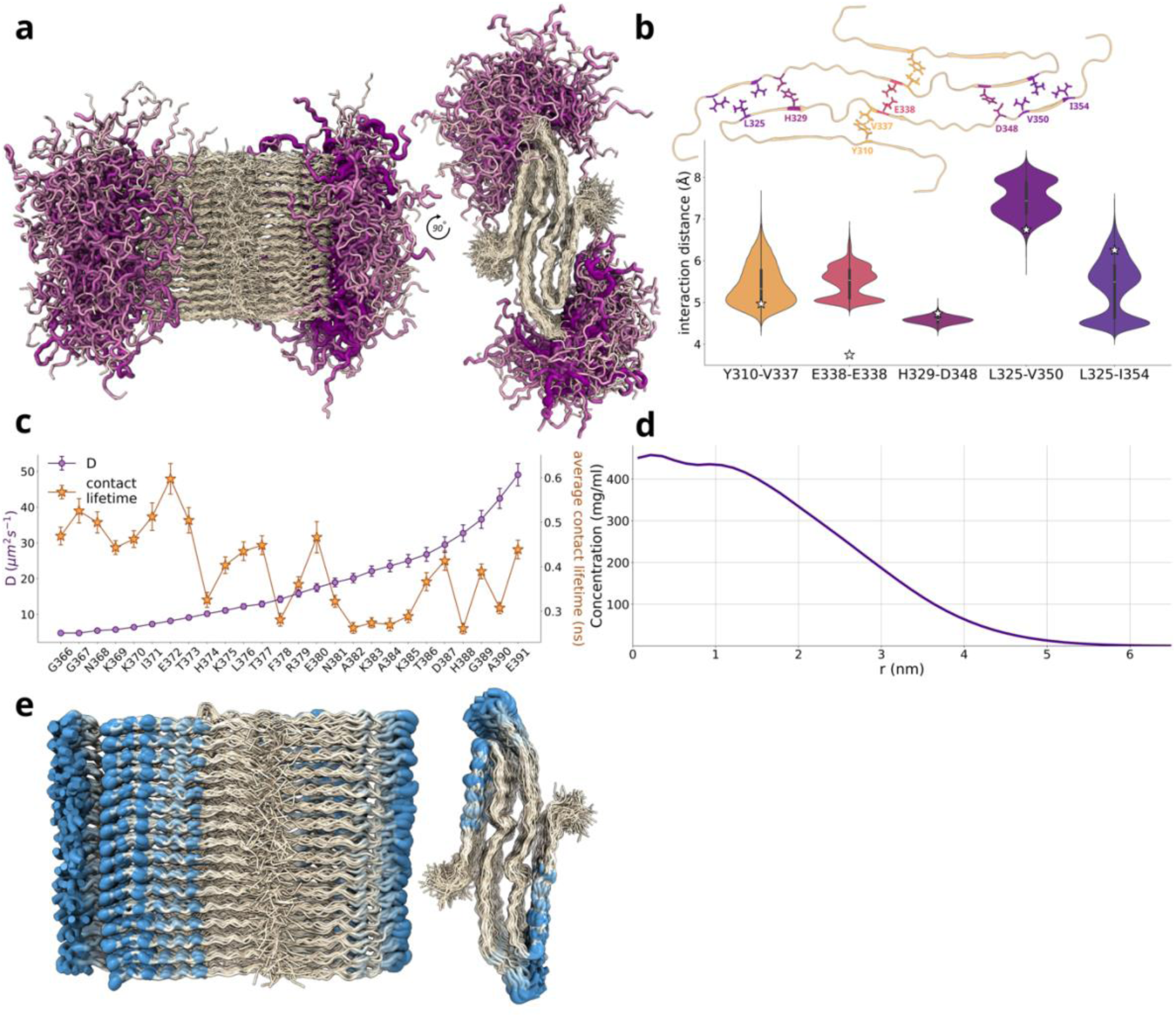
Martini3-NMR reveals the role of the fuzzy coats in the conformational ensembles of a non-twisting tau-amyloid. (a) Ensembles of the 297-391 tau amyloid as sampled by Martini3-NMR. Core residues are coloured white and the residues of the intrinsically disordered regions flanking the core (G366 to E391) are shown using a worm-like representation. The width and the colour intensity of the chain are proportional to the diffusion coefficients shown in panel c. (b) Distance distributions (as violin plots) of the interacting residues deemed as stabilising the tau amyloid formed by residues 297-391.^41^ The stars show the centre of mass distance calculated from the NMR-resolved structure of the fibril (PDB ID: 8G58). (c) Residue-specific diffusion coefficients (indigo) and contact lifetimes (orange) calculated for the IDRs flanking the amyloid core. (d) Radial density profile of the IDRs as a function of the distance from the core. (e) Projection of the number of contacts between the IDRs and the core. For clarity, the intrinsically disordered regions are omitted.

In addition to investigating the properties of this amyloid core, we analysed the characteristics of the flanking disordered regions. Increasing structural, kinetic, and cellular evidence supports the role of these regions, designated as ‘fuzzy coat’^42–45^, as key regulatory interfaces that promote and modulate secondary nucleation as well as tune of intermolecular interactions in ways that strongly influence the toxicity, seeding efficiency, and overall biological behaviour of amyloids. Because current high-resolution structural methods are primarily effective in resolving the ordered fibril core, the conformational properties of disordered fuzzy-coat regions have remained inaccessible, underscoring the need for new approaches capable of elucidating the nature of these heterogeneous and functionally critical regions. When examining the fuzzy coat of the Tau_297–391_ amyloid, the analyses revealed that the pattern of contacts with ordered regions is largely preserved across the disordered sequence (Figure S7b), with only partial reduction of the average contact lifetime in the C-terminal moiety of the sequence compared with the N-terminal residues lying in closer proximity to the structured core. By contrast, a significant difference between residues at the two extremities of the disordered sequence was found when analysing the diffusion coefficient D, with a fivefold increase in D for the C-terminal residues (Figure 5c). Remarkably, the density profile of the fuzzy coat was found to resemble that of protein condensates (Figure 5d). Consistent with behaviour observed in protein condensates,^46,47^ the IDR residues remained highly dynamic despite their elevated local concentration near the fibril core, forming numerous yet transient interactions with the core (Figure 5e) and with other disordered segments. ^46^Overall, by implementing CS and NOE restraints Martini3-NMR was able to recapitulate critical interactions and dynamics in an otherwise challenging system for coarse-grained models.

## Discussion

It is now widely recognised that structural dynamics in proteins are a key modulator of biological activity. Their role is particularly critical for intrinsically disordered proteins, whose interactions, phase-separation behaviour and a wide range of other biomolecular properties depend predominantly on their inherent conformational flexibility. Despite this central relevance, protein dynamics remain challenging to characterise at a molecular level. In this context, CG simulations provide a powerful framework for sampling the conformational space of biomacromolecules, enabling access to timescales and system sizes that are prohibitive for full-atom MD. Such models have greatly expanded our ability to investigate large-scale molecular motions, supramolecular assemblies and long-timescale events that are central to biological function.^48,49^ However, the reduced representation of the system and simplified interaction potentials that underpin CG models introduce intrinsic limitations, often leading to the sampling of conformational states that are unlikely to be significantly populated under physiological conditions.

Here, we demonstrate that artificial neural networks can be employed to develop a methodology for modelling NMR chemical shifts within CG protein force fields, thereby enabling the incorporation of these data as experimental restraints to substantially improve the conformational description of proteins in such simplified representations. This integrative approach broadens the scope of molecular dynamics simulations by combining the enhanced sampling efficiency of CG models with the structural accuracy afforded by NMR restraints. We illustrate the impact of this strategy by generating CG conformational ensembles for a diverse range of systems, including soluble proteins, membrane proteins, amyloid assemblies and intrinsically disordered protein regions.

The application of NapShift-CG to soluble systems, such as a *de novo*–designed mini-protein, yielded excellent agreement with experimental observations. However, for systems featuring more complex topologies, such as ubiquitin, the application of chemical shift restraints within the Martini3 framework primarily improved backbone dihedral sampling, while tertiary contacts were not fully preserved. By contrast, the simultaneous incorporation of NOE and chemicalshift restraints led to a marked improvement in simulation quality, resulting in substantially enhanced representations of both protein structure and dynamics for systems such as Ubiquitin, KRAS and similar proteins. The enhanced capabilities of Martini3-NMR are further illustrated by simulations initiated from a misfolded conformation of ubiquitin, which rapidly folded into the native structure and subsequently explored a conformational ensemble closely aligned with the experimental reference throughout the remainder of the simulation. In contrast, simulations performed using elastic-network Martini3 remained trapped in the misfolded state, while Martini-DSSP produced a heterogeneous set of misfolded, molten-globule–like conformations.

The breadth of applications of Martini3-NMR is further illustrated by its ability to capture the disassembly mechanism of pentameric phospholamban (PLN) within a lipid bilayer. This result demonstrates that complex processes occurring *in situ* in biological membranes, including the organisation and rearrangement of protein assemblies, can be effectively investigated using the proposed approach. Another relevant application of Martini3-NMR concerns the characterisation of the fuzzy regions of amyloid fibrils, which constitute critical sites for their propagation and biological activity. Our results indicate that these regions exhibit spatial organisation, density, and dynamic properties reminiscent of protein condensates, highlighting their potential role as functionally relevant, phase-separation-like environments at the fibril surface.

In conclusion, the methodology introduced here opens new opportunities for extending CG simulations toward a more realistic and experimentally grounded description of protein conformational landscapes. By integrating NMR observables directly into CG force fields, this approach provides a general framework for interrogating dynamic, heterogeneous and multiscale processes that are otherwise difficult to access using existing computational or experimental methodologies in isolation. Martini3-NMR offers a powerful strategy for studying systems in which structural dynamics are central to function, including intrinsically disordered proteins, membrane-associated assemblies, amyloids and phase-separated condensates. Further extensions of this framework to incorporate additional experimental observables and to address increasingly complex cellular environments are expected to enable CG simulations to assume a more quantitative role in linking molecular structure and dynamics to biological function.

## Materials and Methods

### NapShift ANN

We trained a feed-forward Artificial Neural Network (ANN) to predict secondary CS of the 6 backbone atoms (N, C_α_, C_b,_ C, HN and H_α_). A set of 3,236 NMR structures of proteins was gathered from the Protein Data Bank (PDB), with associated CS sourced from the BioMagnetic Resonance Data Bank (BMRB).^50^ Where a given PDB file contained multiple structures, only the first (lowest energy) structure was considered. We filtered out erroneous CS entries by discarding those more than 3 standard deviations from their average value in the BMRB. Raw CS were converted to secondary CS by subtracting their random-coil CS, as calculated by CamCoil.^51^

The structural properties of each residue were analysed using a set of angles describing its local geometry and a vector encoding its amino acid type. While the all-atom ANN CS Predictor^24^ relied on the ϕ, ψ, χ1, and χ2 angles as geometric features, these atomistic resolution observables are unavailable in MARTINI3. We therefore leveraged the angle and dihedrals between a given residue’s backbone atom (BB) and those of the residues preceding and succeeding it in the protein sequence (see angles β, θ_1_ and θ_1_ in figure 1a). We also described the orientation of the residue’s side chain by including the angles involving the first side chain atom (SC1), termed α and γ (Figure 1a). When composing the input vector, each of these angles is represented by their sine and cosine values. Missing angles (e.g. a non-existent θ_1_ for a C-terminal residue) were given values of 0. An embedding of amino acid type was obtained from the BLOSUM62 matrix. This contained entries for the standard 20 amino acids, plus two additional to represent oxidized cysteine (CYO) and cis-proline (PRC).

In this configuration, each MARTINI3 residue is therefore represented by a vector of 32 values (22 encoding amino-acid type, and 5x2 describing its angular geometry). In particular, a residue’s input vector was concatenated with those of the residues before and after in sequence to produce a tri-peptide representation ( vector of 32x3=96 values). The architecture of the ANN involved an input layer (size 96), a hidden layer (size 26) equipped with the Exponential Linear Unit (ELU) activation function^52^, and an output layer (size 6) with a linear activation function, as typical of other regression models. Secondary CS for the 6 backbone atoms were predicted simultaneously.

Collectively, 272,984 tripeptides extracted from 2,687 structures comprised the training set for the NapShift ANN. A Mean Squared Error (MSE) loss function and the ADAM optimizer^53^ with a learning rate of 0.001 were used to train the ANN. After each epoch, an early stopping criterion was assessed on a validation set of 32,076 tripeptides from 299 structures. This served to mitigate overfitting by halting training if the validation loss did not improve after 5 epochs. The best model obtained (according to validation performance) was tested on a set of 28,107 tripeptides extracted from the 250 test structures. Structures were not mixed across the training, validation, and test sets to avoid data leakage.

A harmonic restraint potential is formed by the difference between CS predicted by the ANN for a simulated conformation and those measured by NMR experiments:

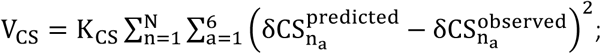

where δCS^predicted^is the secondary CS predicted by the ANN, δCS^observed^is the experimentally observed value, and K_CS_ is the force constant of the restraint potential. The CS restraint potential is slowly ‘ramped up’ by increasing K_CS_ from 0 by 0.001 per step until its maximum value is reached. The CS restraint potential has been implemented as a plugin for the OpenMM simulation engine with GPU support.^25^

### NOE Restraints

In Martini3-NMR, tertiary contacts were accounted through the structural information provided by NOESY experiments. As in the case of the CS restraints, we first translated the interatomic distances associated with NOE data into MARTINI3-compatible distances. In particular, for a given atom pair a_i_, a_j_ with an NOE-derived interatomic distance d_ij_, and coarse-grained into MARTINI beads A_i_, A_j_ respectively, we calculated their CG NOE distance D(A_i_, A_j_) as:

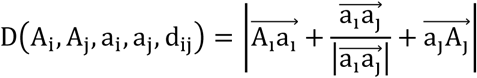

It is common for multiple groups of atoms to contribute to the same NOE observable - for example, in protons of methyl groups, giving rise to a single NOE data. When handling this situation in atomistic-resolution simulations, as in Gromacs 2025,^54^ a single distance restraint is applied across all atoms involved, which calculates its ‘apparent distance’ as:

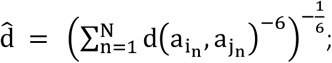

since the NOE signal itself is inversely proportional to the sixth power of the inter-atomic distance. We here employed a similar concept to condense multiple inter-atomic distances into a single CG distance:

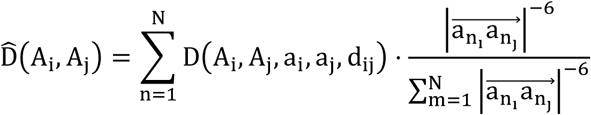

Even though signals from multipair atomistic NOEs are condensed into a single coarse-grain distance, there may still be multiple different experimental signals which map onto the same MARTINI3 bead pair, A_i_ A_j_ – for example, one might measure H_αi_ -H_αj_ and C_αi_-H_αj_ , corresponding to two separate interactions between BB_i_ and BB_j_ . In this case, the resultant coarse-grain distance is calculated as the average coarse-grain distance for that MARTINI bead pair:

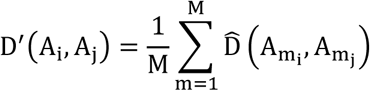

Finally, we considered the case where more than one residue pair contributes to an NOE signal. In such cases, we filtered these by checking conflicting residue pairs and discarding those with an inter-atomic distance significantly larger than that reported by the NOE.

Having obtained CG NOE distances, we apply the potential proposed by Torda and van Gusteren:^55^

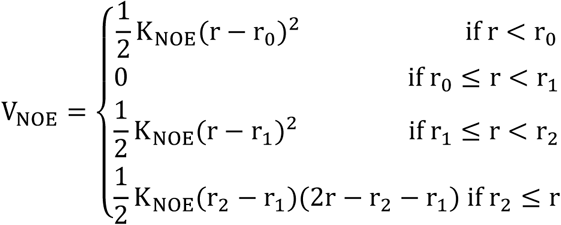

where r is the distance between two MARTINI3 beads and K_NOE_ is the force constant of the NOE restraint potential. This potential includes a harmonic regime, a flat-bottom between r_0_ and r_1_ to account for uncertainty in the measurement, and a linear regime beyond r2. Atomistic r_0_, r_1_ and r_2_ can usually be read directly from NOE data files, but where r_0_ and r_2_are missing, we set r_0_ = 0 (to avoid imposing undue bias on the system) and r_2_ = r_1_ + 0.5 nm.

We also considered the possibility that NOSEY results may imply distance restraints which are unable to be satisfied instantaneously, corresponding to different structural states experienced by the system. To account for this, we applied NOE restraints to time-averaged distances instead of instantaneous distances, as in Torda et al.^56^, resulting in a restraining force of:

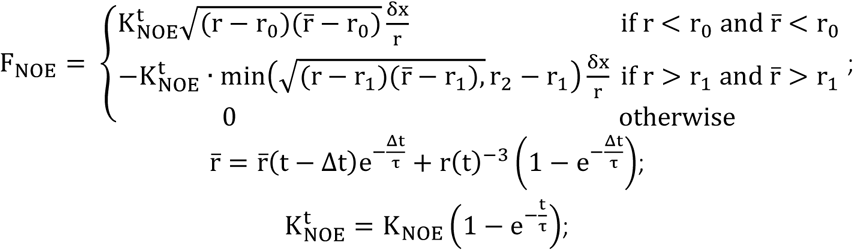

where δx is the direction vector between the two C_α_ beads, t is the current simulation step, and τ = 0.05 ps is the decay time for the exponential running average. As with the CS restraint potential, this NOE restraint potential has been implemented as a plugin for OpenMM. The complete set of NMR restraint potentials (CS and NOE) are added to the potential energy arising from the MARTINI forcefield to produce a total potential energy of:

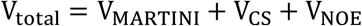

*Data Preparation:* The martinize2 script^28^ version 0.13.0 was used to map experimental protein structures to their MARTINI3 representation. The cys –auto flag was used to detect disulfide bonds within systems. Membrane systems were generated from atomistic protein structures using CHARMM-

GUI’s martini-maker tool.^57^ For each protein simulated, two MARTINI3 topologies were generated: one without MARTINI3 secondary structure or scFix^58^ restraints for simulation with NMR restraints, and one with MARTINI3 restraints for secondary structure detected by DSSP^59^ and scFix applied for simulation without NMR restraints. Having mapped structures according to the MARTINI3 mapping scheme, GROMACS 2024.5^60^ was used to place the CG models at the centre of a cubic box such that all protein atoms were at least 1.2 nm from the box boundaries. The system was then solvated with MARTINI3 water beads, after which NA^+^ CL^-^ ions were added with gmx addion, bringing the system to the desired salt concentration, as reported in the excel spreadsheet in the supplementary material.

Parameters for NOE restraints were computed using experimental data reported and structures sourced from the PDB. CS were acquired from the corresponding BMRB entries. Where specified, the full-atom NapShift ^24^ was used to generate synthetic CS from the full atom experimental structure in cases where the experimental CS dataset was not available or largely incomplete.

To assess the ability of Martini3-NMR to recover native-like conformations, we generated an atomistic partially unfolded conformation for all-atom Ubiquitin as shown in Figure 2, via simulated annealing using the CHARMM36 force field^61^. After energy minimization, the system was equilibrated in the NVT and NPT ensembles, before a simulated annealing run of 5 ‘cycles’ was performed: for each cycle, the simulation temperature was kept constant at 300K for 5 ns, gradually raised to 800K over 2 ns, kept constant at 800K for a further 5 ns, before being lowered back to 300K over 2 ns. The final frame of this simulation was used as the ‘melted’ initial atomistic structure.

The structure of the AFA Phospholamban mutant was obtained by mutating Cys36Ala, Cys41Phe, and Cys46Ala in each transmembrane helix of the atomistic structure in PDB ID 2KYV with ChimeraX.^62^ The same set of experimental NOEs (provided in PDB 2KYV) were used for WT and AFA Phospholamban, although their mapping and the prediction of synthetic CS were performed independently for the two initial structures. For simulations investigating the dissociation of transmembrane helix 1, all intermolecular NOEs involving transmembrane helix 1 were removed.

The structure of the Tau amyloid was taken from the 5-layer atomistic structure resolved by ssNMR (PDB ID 8G58). This structure was duplicated and translated 4 times to create a 20-layer fiber. NOE and CS values were also duplicated to create corresponding restraints for the generated layers. CS for this system were complemented with those generated synthetically with NapShift ^24^ and CamCoil ^51^ for the disordered regions. Initial structures for the disordered tails were modelled as extended conformations using the [X] tool provided by ChimeraX.^62^

*Simulations:* Coarse-grained Langevin dynamics simulations were carried by OpenMM 8.2.0,^25^ using the Langevin integrator with a time step of 10 fs and friction coefficient of 0.01 ps^-1^, under periodic boundary conditions. All systems were simulated under the MARTINI3 forcefield.^63^ Gromacs topology files defining MARTINI3 systems were converted into OpenMM systems by the martini-openmm tool.^64^ At the beginning of each simulation, the system was energy-minimized by the steepest descent algorithm, followed by 500 ps of each NVT and NPT equilibration, during which protein atoms were positionally restrained. The CS restraints were then gradually imposed by increasing K_CS_ by 0.001 each step until the pre-determined maximum value (reported for each system in Supplementary data spreadsheet) was reached. All systems were simulated in replicates of 3. Total equilibration and simulation times are reported in excel spreadsheet provided in the supplementary data. Convergence of simulation trajectories was assessed by applying the Augmented Dickey Fuller (ADF) test to all timeseries data presented. ADF test statistics and p-values are reported in Supplementary data spreadsheet.

*Analysis:* <u>Backbone Cα RMSD</u> values (Figures 1c, 2a, 2b, 3a, 3b) were calculated by first aligning all structures of an ensemble to the starting structure, then computing the RMSD using the BB beads in each simulation frame. Only structured regions were considered in the alignment and calculation of backbone RMSD. For the two-domain system 9COJ shown in Figure 3a, the alignment procedure was repeated considering each of the two domains separately. For the system 2KYV, backbone RMSD was calculated considering only the helical transmembrane domain.

Dihedral RMSD values (Figures 2c, 3a-b) were similarly computed by taking the RMSD of each backbone dihedral angle with respect to the starting structure of the simulations.

<u>CS agreement</u> (Figure 1c) was computer per-residue by comparing experimental and calculated CS for simulations with and without NMR restraints (Figure S1) and compared via their Pearson correlation coefficients. For a given conformation, the agreement between simulated and experimental CS was condensed to a single number by computing the RMSD of simulated from experimental CS for each atom type and scaling this according to the standard deviation for CS of this atom type observed in the BMRB.

Pseudo-Ramachandran plots (Figures S2c, S5) were obtained by calculating and plotting the main chain dihedral angles θ_1_𝑎𝑛𝑑 θ_2_ (Figure 1a) for each residue of the protein and across the ensemble structures.

<u>τ and ω angles</u> (Figure 4) were computed as in Sanz-Hernandez *et al*. ^38^ using the vectors B_0_ (the membrane normal), h⃗ (the direction vector of a single transmembrane helix i), and h⃗ (the direction vector of the entire transmembrane helical bundle). h⃗ was calculated by taking the difference between the positions of the backbone beads at the beginning and end of transmembrane helix i (start_TMi_ and end_TMi_), and h⃗ _bundle_ was calculated as:

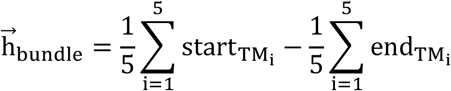

and subsequently compute:

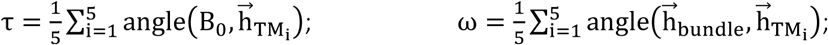

<u>The dissociation of transmembrane helix 1 from the phospholamban pentamer</u> was calculated using the total number of contacts made by the helix with the rest of the transmembrane bundle using a cutoff of 0.8 nm to assign a contact. A conformation was considered ‘dissociated’ if the number of contacts was found to be 0. Histograms of contact counts were transformed into free energy surfaces by calculating:

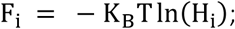

where 𝐹_𝑖_ is the free energy calculated for histogram bin 𝑖 , 𝐾_𝐵_ is the Boltzmann constant, 𝑇 is the simulation temperature, and 𝐻_𝑖_ is the density in bin 𝑖 . Errors on 𝐹 were calculated as:

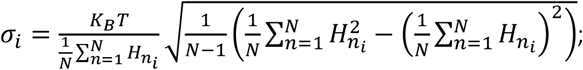

where 𝐻_𝑛_ is the n^th^ histogram of 𝑁 bootstrap samples.

<u>In the analysis of the tau-amyloid</u>, the first and last two layers of the amyloid assembly (totalling 20 layers encompassing 40 chains in our model) were discarded to avoid end-effects in the fibril. Mean-square displacement (MSD) of the amyloid’s disordered tails was computed by first aligning all frames of an amyloid ensemble using the backbone beads of the structured fibril core. The MSD of a given backbone bead in a disordered tail was calculated using MDAnalysis^65^ with a minimum lag time of 10 ps (the reporting frequency of the simulation). We consider the segment between 100ps and 500ps to correspond to the linear regime of MSD. For each residue, its diffusion coefficient was computed by fitting a linear model to this linear regime. Pearson correlation coefficients for these fits are reported in the Supplementary data spreadsheet. Tail residues in the ensemble shown in Figure 5c are colored according to the natural log of their computed 𝐷.

Contact lifetime analysis employed the transition-based definition described in Galvanetto et al.^46^, whereby a contact between two residues is considered formed when the minimum distance of all bead-pairs for these two residues drops below r_0_ and considered broken the next time that this minimum distance rises above r_1_. We set r_0_=0.8 nm, and r_1_=1.0 nm. The average contact lifetime of a given tail residue with the fiber core was calculated by taking the average of residue-residue contact lifetimes across all 36 chains, then for each tail residue taking its average contact lifetime with all core residues. Contacts that were not formed in any replicate simulation were not included in this average.

The radial concentration profile of the disordered tau amyloid tails was computed as:

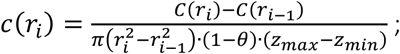

where 𝐶(𝑟_𝑖_) is the number of tail atoms less than 𝑟_𝑖_ nm from the point of attachment to the fiber core in the xy plane, 𝜃 = ^𝜋/2^ is the proportion of the circular area occupied by the structured core (which should be excluded from the density calculation), and 𝑧_𝑚𝑖𝑛_, 𝑧_𝑚𝑎𝑥_ are the z-coordinates of the bottom and top of the core respectively. Tail atoms with z coordinates higher than 𝑧_𝑚𝑎𝑥_ or lower than 𝑧_𝑚𝑖𝑛_ were not included in the calculation.

The twist angle between successive layers of the fiber core was computed as the dihedral between the BB beads L325_a_, L325_b_, L325_c_, L325_d_ in chains a, b (layer 𝑖), c, d (layer 𝑖 + 1), similar to the procedure described in Periole et al.^66^. For each conformation, the average twist angle was calculated and used to produce the distributions shown in Supplementary Figure S7.

*Visualization:* Simulated conformations were visualized using VMD version 1.9.4^66^ and ChimeraX version 1.8^62^. Coarse-grained ensembles (grey lines) in Figures 1c, 2b and 3 are shown paired with a representative structure. These representative structures were obtained by performing a PCA on the ensemble coordinates and selecting the structure closest to the average of the PCA-transformed ensemble.

## Author contributions

M.C. trained the ANN, implemented the framework within OpenMM, ran simulations and analysed data; A.D.S. and D.M. conceived and designed the study; D.M. and K.T. jointly supervised M.C.; C.B., V.M. and A.D.S. advised M.C. on several aspects of the work; A.D.S. wrote the manuscript with the contribution of all the authors.

## Data and code availability

The excel spreadsheet named “SupplementalData.xlsx” provides the following data: The spreadsheet named “Dataset” provides the list of PDB files for the train, validation and test sets. The spreadsheet named “Systems” contains simulation conditions and DOIs for the systems simulated (membrane and soluble proteins). The spreadsheet named “ADF Statistics” provides tables of the Augmented Dickey-Fuller (ADF) coefficients calculated to assess stationarity of all the presented timeseries. Cells highlighted in red report stationarity, within a 5% confidence level, otherwise within a 1% confidence level. The spreadsheet named “MSD Correlation” provides the correlation coefficients obtained from the fit of the linear regime of the mean square displacement traces obtained for each residue shown in Figure 5c. Napshift plugins for OpenMM and Martini3-NMR tutorials will be made available upon publication of the peer-reviewed article.

## Acknowledgements

We are grateful to Prof. Gianluigi Veglia for discussions. This research was supported by the European Research Council (ERC) project BioDisOrder - 819644 to A.D. and Italian Association for Cancer Research (AIRC) project IG 30556.

## References

(1) Holehouse, A. S.; Kragelund, B. B. The Molecular Basis for Cellular Function of Intrinsically Disordered Protein Regions. Nature Reviews Molecular Cell Biology 2023 25:3 2023, 25 (3), 187–211.

(2) Eisenmesser, E. Z.; Bosco, D. A.; Akke, M.; Kern, D. Enzyme Dynamics during Catalysis. Science (1979) 2002, 295 (5559), 1520–1523.

(3) Karamanos, T. K.; Jackson, M. P.; Calabrese, A. N.; Goodchild, S. C.; Cawood, E. E.; Thompson, G. S.; Kalverda, A. P.; Hewitt, E. W.; Radford, S. E. Structural Mapping of Oligomeric Intermediates in an Amyloid Assembly Pathway. Elife 2019, 8.

(4) Wong, W.; Gough, N. R. Focus Issue: The Protein Dynamics of Cell Signaling. Sci Signal 2009, 2 (66).

(5) Neudecker, P.; Robustelli, P.; Cavalli, A.; Walsh, P.; Lundström, P.; Zarrine-Afsar, A.; Sharpe, S.; Vendruscolo, M.; Kay, L. E. Structure of an Intermediate State in Protein Folding and Aggregation. Science (1979) 2012, 336 (6079), 362–366.

(6) Sedinkin, S. L.; Burns, D.; Shukla, D.; Potoyan, D. A.; Venditti, V. Solution Structure Ensembles of the Open and Closed Forms of the ∼130 KDa Enzyme I via AlphaFold Modeling, Coarse Grained Simulations, and NMR. J Am Chem Soc 2023, 145 (24), 13347–13356.

(7) Sučec, I.; Xia, B.; Somberg, N. H.; Wang, Y.; Jo, H.; Li, S.; Perrone, B.; Gao, Z.; Hong, M. Ion Channel Structure and Function of the MERS Coronavirus E Protein. Science Advances 2025, 11 (28), 1788.

(8) Weber, D. K.; Reddy, U. V.; Wang, S.; Larsen, E. K.; Gopinath, T.; Gustavsson, M. B.; Cornea, R. L.; Thomas, D. D.; De Simone, A.; Veglia, G. Structural Basis for Allosteric Control of the SERCA-Phospholamban Membrane Complex by Ca2+ and Phosphorylation. Elife 2021, 10.

(9) Lee, M.; Wang, T.; Makhlynets, O. V.; Wu, Y.; Polizzi, N. F.; Wu, H.; Gosavi, P. M.; Stöhr, J.; Korendovych, I. V.; Degrado, W. F.; Hong, M. Zinc-Binding Structure of a Catalytic Amyloid from Solid-State NMR. Proc Natl Acad Sci U S A 2017, 114 (24), 6191–6196.

(10) Sahoo, B. R.; Genjo, T.; Bekier, M.; Cox, S. J.; Stoddard, A. K.; Ivanova, M.; Yasuhara, K.; Fierke, C. A.; Wang, Y.; Ramamoorthy, A. Alzheimer’s Amyloid-Beta Intermediates Generated Using Polymer-Nanodiscs. Chemical Communications 2018, 54 (91), 12883–12886.

(11) Tycko, R. The Evolving Role of Solid State Nuclear Magnetic Resonance Methods in Studies of Amyloid Fibrils. Curr Opin Struct Biol 2025, 92, 103043.

(12) Ahmed, R.; Akcan, M.; Khondker, A.; Rheinstädter, M. C.; Bozelli, J. C.; Epand, R. M.; Huynh, V.; Wylie, R. G.; Boulton, S.; Huang, J.; Verschoor, C. P.; Melacini, G. Atomic Resolution Map of the Soluble Amyloid Beta Assembly Toxic Surfaces. Chem Sci 2019, 10 (24), 6072–6082.

(13) Mainz, A.; Religa, T. L.; Sprangers, R.; Linser, R.; Kay, L. E.; Reif, B. NMR Spectroscopy of Soluble Protein Complexes at One Mega-Dalton and Beyond. Angewandte Chemie International Edition 2013, 52 (33), 8746–8751.

(14) Sprangers, R.; Kay, L. E. Quantitative Dynamics and Binding Studies of the 20S Proteasome by NMR. Nature 2006 445:7128 2007, 445 (7128), 618–622.

(15) Murray, D. T.; Kato, M.; Lin, Y.; Thurber, K. R.; Hung, I.; McKnight, S. L.; Tycko, R. Structure of FUS Protein Fibrils and Its Relevance to Self-Assembly and Phase Separation of Low-Complexity Domains. Cell 2017, 171 (3), 615–627.e16.

(16) Wake, N.; Weng, S. L.; Zheng, T.; Wang, S. H.; Kirilenko, V.; Mittal, J.; Fawzi, N. L. Expanding the Molecular Grammar of Polar Residues and Arginine in FUS Phase Separation. Nature Chemical Biology 2025 21:7 2025, 21 (7), 1076–1088.

(17) De Simone, A.; Richter, B.; Salvatella, X.; Vendruscolo, M. Toward an Accurate Determination of Free Energy Landscapes in Solution States of Proteins. J Am Chem Soc 2009, 131 (11), 3810–3811.

(18) Montalvao, R. W.; De Simone, A.; Vendruscolo, M. Determination of Structural Fluctuations of Proteins from Structure-Based Calculations of Residual Dipolar Couplings. J Biomol NMR 2012, 53 (4), 281–292.

(19) Camilloni, C.; Robustelli, P.; Simone, A. De; Cavalli, A.; Vendruscolo, M. Characterization of the Conformational Equilibrium between the Two Major Substates of RNase a Using NMR Chemical Shifts. J Am Chem Soc 2012, 134 (9), 3968–3971.

(20) De Simone, A.; Aprile, F. A.; Dhulesia, A.; Dobson, C. M.; Vendruscolo, M. Structure of a Low-Population Intermediate State in the Release of an Enzyme Product. Elife 2015, 2015 (4).

(21) De Simone, A.; Montalvao, R. W.; Dobson, C. M.; Vendruscolo, M. Characterization of the Interdomain Motions in Hen Lysozyme Using Residual Dipolar Couplings as Replica-Averaged Structural Restraints in Molecular Dynamics Simulations. Biochemistry 2013, 52 (37), 6480–6486.

(22) De Simone, A.; Montalvao, R. W.; Vendruscolo, M. Determination of Conformational Equilibria in Proteins Using Residual Dipolar Couplings. J Chem Theory Comput 2011, 7 (12), 4189–4195.

(23) De Simone, A.; Gustavsson, M.; Montalvao, R. W.; Shi, L.; Veglia, G.; Vendruscolo, M. Structures of the Excited States of Phospholamban and Shifts in Their Populations upon Phosphorylation. Biochemistry 2013, 52 (38), 6684–6694.

(24) Qi, G.; Vrettas, M. D.; Biancaniello, C.; Sanz-Hernandez, M.; Cafolla, C. T.; Morgan, J. W. R.; Wang, Y.; De Simone, A.; Wales, D. J. Enhancing Biomolecular Simulations with Hybrid Potentials Incorporating NMR Data. J Chem Theory Comput 2022, 18 (12), 7733–7750.

(25) Eastman, P.; Galvelis, R.; Peláez, R. P.; Abreu, C. R. A.; Farr, S. E.; Gallicchio, E.; Gorenko, A.; Henry, M. M.; Hu, F.; Huang, J.; Krämer, A.; Michel, J.; Mitchell, J. A.; Pande, V. S.; Rodrigues, J. P.; Rodriguez-Guerra, J.; Simmonett, A. C.; Singh, S.; Swails, J.; Turner, P.; Wang, Y.; Zhang, I.; Chodera, J. D.; De Fabritiis, G.; Markland, T. E. OpenMM 8: Molecular Dynamics Simulation with Machine Learning Potentials. Journal of Physical Chemistry B 2024, 128 (1), 109–116.

(26) Bhardwaj, G.; Mulligan, V. K.; Bahl, C. D.; Gilmore, J. M.; Harvey, P. J.; Cheneval, O.; Buchko, G. W.; Pulavarti, S. V. S. R. K.; Kaas, Q.; Eletsky, A.; Huang, P. S.; Johnsen, W. A.; Greisen, P. J.; Rocklin, G. J.; Song, Y.; Linsky, T. W.; Watkins, A.; Rettie, S. A.; Xu, X.; Carter, L. P.; Bonneau, R.; Olson, J. M.; Coutsias, E.; Correnti, C. E.; Szyperski, T.; Craik, D. J.; Baker, D. Accurate de Novo Design of Hyperstable Constrained Peptides. Nature 2016 538:7625 2016, 538 (7625), 329–335.

(27) Vijay-kumar, S.; Bugg, C. E.; Cook, W. J. Structure of Ubiquitin Refined at 1.8Åresolution. J Mol Biol 1987, 194 (3), 531–544.

(28) Kroon, P. C.; Grünewald, F.; Barnoud, J.; Tilburg, M. van; Brasnett, C.; Souza, P. C.; Wassenaar, T. A.; Marrink, S. J. Martinize2 and Vermouth Provide a Unified Framework for Molecular Topology Generation. Elife 2025, 12, RP90627.

(29) Bocharov, E. V.; Lesovoy, D. M.; Pavlov, K. V.; Pustovalova, Y. E.; Bocharova, O. V.; Arseniev, A. S. Alternative Packing of EGFR Transmembrane Domain Suggests That Protein–Lipid Interactions Underlie Signal Conduction across Membrane. Biochimica et Biophysica Acta (BBA) - Biomembranes 2016, 1858 (6), 1254–1261.

(30) Zanetti-Domingues, L. C.; Korovesis, D.; Needham, S. R.; Tynan, C. J.; Sagawa, S.; Roberts, S. K.; Kuzmanic, A.; Ortiz-Zapater, E.; Jain, P.; Roovers, R. C.; Lajevardipour, A.; van Bergen en Henegouwen, P. M. P.; Santis, G.; Clayton, A. H. A.; Clarke, D. T.; Gervasio, F. L.; Shan, Y.; Shaw, D. E.; Rolfe, D. J.; Parker, P. J.; Martin-Fernandez, M. L. The Architecture of EGFR’s Basal Complexes Reveals Autoinhibition Mechanisms in Dimers and Oligomers. Nature Communications 2018 9:1 2018, 9 (1), 4325-.

(31) Thomasen, F. E.; Skaalum, T.; Kumar, A.; Srinivasan, S.; Vanni, S.; Lindorff-Larsen, K. Rescaling Protein-Protein Interactions Improves Martini 3 for Flexible Proteins in Solution. Nature Communications 2024 15:1 2024, 15 (1), 6645-.

(32) Artemenko, E. O.; Egorova, N. S.; Arseniev, A. S.; Feofanov, A. V. Transmembrane Domain of EphA1 Receptor Forms Dimers in Membrane-like Environment. Biochimica et Biophysica Acta (BBA) - Biomembranes 2008, 1778 (10), 2361–2367.

(33) Shahid, S. A.; Bardiaux, B.; Franks, W. T.; Krabben, L.; Habeck, M.; Van Rossum, B. J.; Linke, D. Membrane-Protein Structure Determination by Solid-State NMR Spectroscopy of Microcrystals. Nature Methods 2012 9:12 2012, 9 (12), 1212–1217.

(34) Verardi, R.; Shi, L.; Traaseth, N. J.; Walsh, N.; Veglia, G. Structural Topology of Phospholamban Pentamer in Lipid Bilayers by a Hybrid Solution and Solid-State NMR Method. Proc Natl Acad Sci U S A 2011, 108 (22), 9101–9106.

(35) Toyoshima, C.; Asahi, M.; Sugita, Y.; Khanna, R.; Tsuda, T.; MacLennan, D. H. Modeling of the Inhibitory Interaction of Phospholamban with the Ca2+ ATPase. Proc Natl Acad Sci U S A 2003, 100 (2), 467–472.

(36) Mravic, M.; Thomaston, J. L.; Tucker, M.; Solomon, P. E.; Liu, L.; DeGrado, W. F. Packing of Apolar Side Chains Enables Accurate Design of Highly Stable Membrane Proteins. Science (1979) 2019, 363 (6434), 1418–1423.

(37) Liu, W.; Fei, J. Z.; Kawakami, T.; Smith, S. O. Structural Constraints on the Transmembrane and Juxtamembrane Regions of the Phospholamban Pentamer in Membrane Bilayers: Gln29 and Leu52. Biochim Biophys Acta 2007, 1768 (12), 2971.

(38) Sanz-Hernández, M.; Vostrikov, V. V.; Veglia, G.; De Simone, A. Accurate Determination of Conformational Transitions in Oligomeric Membrane Proteins. Scientific Reports 2016 6:1 2016, 6 (1), 23063-.

(39) Funk, F.; Kronenbitter, A.; Hackert, K.; Oebbeke, M.; Klebe, G.; Barth, M.; Koch, D.; Schmitt, J. P. Phospholamban Pentamerization Increases Sensitivity and Dynamic Range of Cardiac Relaxation. Cardiovasc Res 2023, 119 (7), 1568–1582.

(40) Weber, D. K.; Reddy, U. V.; Robia, S. L.; Veglia, G. Pathological Mutations in the Phospholamban Cytoplasmic Region Affect Its Topology and Dynamics Modulating the Extent of SERCA Inhibition. Biochimica et Biophysica Acta (BBA) - Biomembranes 2024, 1866 (7), 184370.

(41) Duan, P.; Dregni, A. J.; Mammeri, N. El; Hong, M. Structure of the Nonhelical Filament of the Alzheimer’s Disease Tau Core. Proc Natl Acad Sci U S A 2023, 120 (44), e2310067120.

(42) Milanesi, M.; Brotzakis, Z. F.; Vendruscolo, M. Transient Interactions between the Fuzzy Coat and the Cross-β Core of Brain-Derived Aβ42 Filaments. Science Advances 2025, 11 (3), 7008.

(43) Ulamec, S. M.; Brockwell, D. J.; Radford, S. E. Looking Beyond the Core: The Role of Flanking Regions in the Aggregation of Amyloidogenic Peptides and Proteins. Front Neurosci 2020, 14, 611285.

(44) Faidon Brotzakis, Z.; Löhr, T.; Truong, S.; Hoff, S.; Bonomi, M.; Vendruscolo, M. Determination of the Structure and Dynamics of the Fuzzy Coat of an Amyloid Fibril of IAPP Using Cryo-Electron Microscopy. Biochemistry 2023, 62 (16), 2407–2416.

(45) Wegmann, S.; Medalsy, I. D.; Mandelkow, E.; Müller, D. J. The Fuzzy Coat of Pathological Human Tau Fibrils Is a Two-Layered Polyelectrolyte Brush. Proc Natl Acad Sci U S A 2012, 110 (4), E313–E321.

(46) Galvanetto, N.; Ivanović, M. T.; Chowdhury, A.; Sottini, A.; Nüesch, M. F.; Nettels, D.; Best, R. B.; Schuler, B. Extreme Dynamics in a Biomolecular Condensate. Nature 2023 619:7971 2023, 619 (7971), 876–883.

(47) Ivanović, M. T.; Best, R. B. All-Atom Simulations of Biomolecular Condensates. Curr Opin Struct Biol 2025, 93, 103101.

(48) Pipatpadungsin, N.; Chao, K.; Rouse, S. L. Coarse-Grained Simulations of Adeno-Associated Virus and Its Receptor Reveal Influences on Membrane Lipid Organization and Curvature. Journal of Physical Chemistry B 2024.

(49) Jefferies, D.; Shearer, J.; Khalid, S. Role of O-Antigen in Response to Mechanical Stress of the E. Coli Outer Membrane: Insights from Coarse-Grained MD Simulations. Journal of Physical Chemistry B 2019, 123 (17), 3567–3575.

(50) Ulrich, E. L.; Akutsu, H.; Doreleijers, J. F.; Harano, Y.; Ioannidis, Y. E.; Lin, J.; Livny, M.; Mading, S.; Maziuk, D.; Miller, Z.; Nakatani, E.; Schulte, C. F.; Tolmie, D. E.; Kent Wenger, R.; Yao, H.; Markley, J. L. BioMagResBank. Nucleic Acids Res 2008, 36 (suppl_1), D402–D408.

(51) De Simone, A.; Cavalli, A.; Hsu, S. T. D.; Vranken, W.; Vendruscolo, M. Accurate Random Coil Chemical Shifts from an Analysis of Loop Regions in Native States of Proteins. J Am Chem Soc 2009, 131 (45), 16332–16333.

(52) Clevert, D. A.; Unterthiner, T.; Hochreiter, S. Fast and Accurate Deep Network Learning by Exponential Linear Units (ELUs). 4th International Conference on Learning Representations, ICLR 2016 - Conference Track Proceedings 2015.

(53) Kingma, D. P.; Ba, J. L. Adam: A Method for Stochastic Optimization. 3rd International Conference on Learning Representations, ICLR 2015 - Conference Track Proceedings 2014.

(54) Abraham, M.; Alekseenko, A.; Andrews, B.; Basov, V.; Bauer, P.; Bird, H.; Briand, E.; Brown, A.; Doijade, M.; Fiorin, G.; Fleischmann, S.; Gorelov, S.; Gouaillardet, G.; Gray, A.; Irrgang, M. E.; Jalalypour, F.; Johansson, P.; Kutzner, C.; Łazarski, G.; Lemkul, J. A.; Lundborg, M.; Merz, P.; Miletić, V.; Morozov, D.; Müllender, L.; Nabet, J.; Páll, S.; Pasquadibisceglie, A.; Pellegrino, M.; Piasentin, N.; Rapetti, D.; Sadiq, M. U.; Santuz, H.; Schulz, R.; Shirts, M.; Shugaeva, T.; Shvetsov, A.; Turner, P.; Villa, A.; Wingbermühle, S.; Hess, B.; Lindahl, E. GROMACS 2025.1 Manual.

(55) Torda, A. E.; van Gunsteren, W. F. The Refinement of NMR Structures by Molecular Dynamics Simulation. Comput Phys Commun 1991, 62 (2–3), 289–296.

(56) Torda, A. E.; Scheek, R. M.; van Gunsteren, W. F. Time-Dependent Distance Restraints in Molecular Dynamics Simulations. Chem Phys Lett 1989, 157 (4), 289–294.

(57) Qi, Y.; Ingólfsson, H. I.; Cheng, X.; Lee, J.; Marrink, S. J.; Im, W. CHARMM-GUI Martini Maker for Coarse-Grained Simulations with the Martini Force Field. J Chem Theory Comput 2015, 11 (9), 4486–4494.

(58) Herzog, F. A.; Braun, L.; Schoen, I.; Vogel, V. Improved Side Chain Dynamics in MARTINI Simulations of Protein-Lipid Interfaces. J Chem Theory Comput 2016, 12 (5), 2446–2458.

(59) Kabsch, W.; Sander, C. Dictionary of Protein Secondary Structure: Pattern Recognition of Hydrogen-Bonded and Geometrical Features. Biopolymers 1983, 22 (12), 2577–2637.

(60) Berendsen, H. J. C.; van der Spoel, D.; van Drunen, R. GROMACS: A Message-Passing Parallel Molecular Dynamics Implementation. Comput Phys Commun 1995, 91 (1–3), 43–56.

(61) Best, R. B.; Zhu, X.; Shim, J.; Lopes, P. E. M.; Mittal, J.; Feig, M.; MacKerell, A. D. Optimization of the Additive CHARMM All-Atom Protein Force Field Targeting Improved Sampling of the Backbone φ, ψ and Side-Chain Χ1 and Χ2 Dihedral Angles. J Chem Theory Comput 2012, 8 (9), 3257–3273.

(62) Meng, E. C.; Goddard, T. D.; Pettersen, E. F.; Couch, G. S.; Pearson, Z. J.; Morris, J. H.; Ferrin, T. E. UCSF ChimeraX: Tools for Structure Building and Analysis. Protein Science 2023, 32 (11), e4792.

(63) Souza, P. C. T.; Alessandri, R.; Barnoud, J.; Thallmair, S.; Faustino, I.; Grünewald, F.; Patmanidis, I.; Abdizadeh, H.; Bruininks, B. M. H.; Wassenaar, T. A.; Kroon, P. C.; Melcr, J.; Nieto, V.; Corradi, V.; Khan, H. M.; Domański, J.; Javanainen, M.; Martinez-Seara, H.; Reuter, N.; Best, R. B.; Vattulainen, I.; Monticelli, L.; Periole, X.; Tieleman, D. P.; de Vries, A. H.; Marrink, S. J. Martini 3: A General Purpose Force Field for Coarse-Grained Molecular Dynamics. Nature Methods 2021 18:4 2021, 18 (4), 382–388.

(64) MacCallum, J. L.; Hu, S.; Lenz, S.; Souza, P. C. T.; Corradi, V.; Tieleman, D. P. An Implementation of the Martini Coarse-Grained Force Field in OpenMM. Biophys J 2023, 122 (14), 2864–2870.

(65) Michaud-Agrawal, N.; Denning, E. J.; Woolf, T. B.; Beckstein, O. MDAnalysis: A Toolkit for the Analysis of Molecular Dynamics Simulations. J Comput Chem 2011, 32 (10), 2319–2327.

(66) Humphrey, W.; Dalke, A.; Schulten, K. VMD: Visual Molecular Dynamics. J Mol Graph 1996, 14 (1), 33–38.

