## Supplementary Information for "Integrating NMR restraints into coarse-grained simulations: toward accurate conformational ensembles of complex protein systems"

**This PDF file includes:**

Figures S1 to S7

### Supplementary Figures

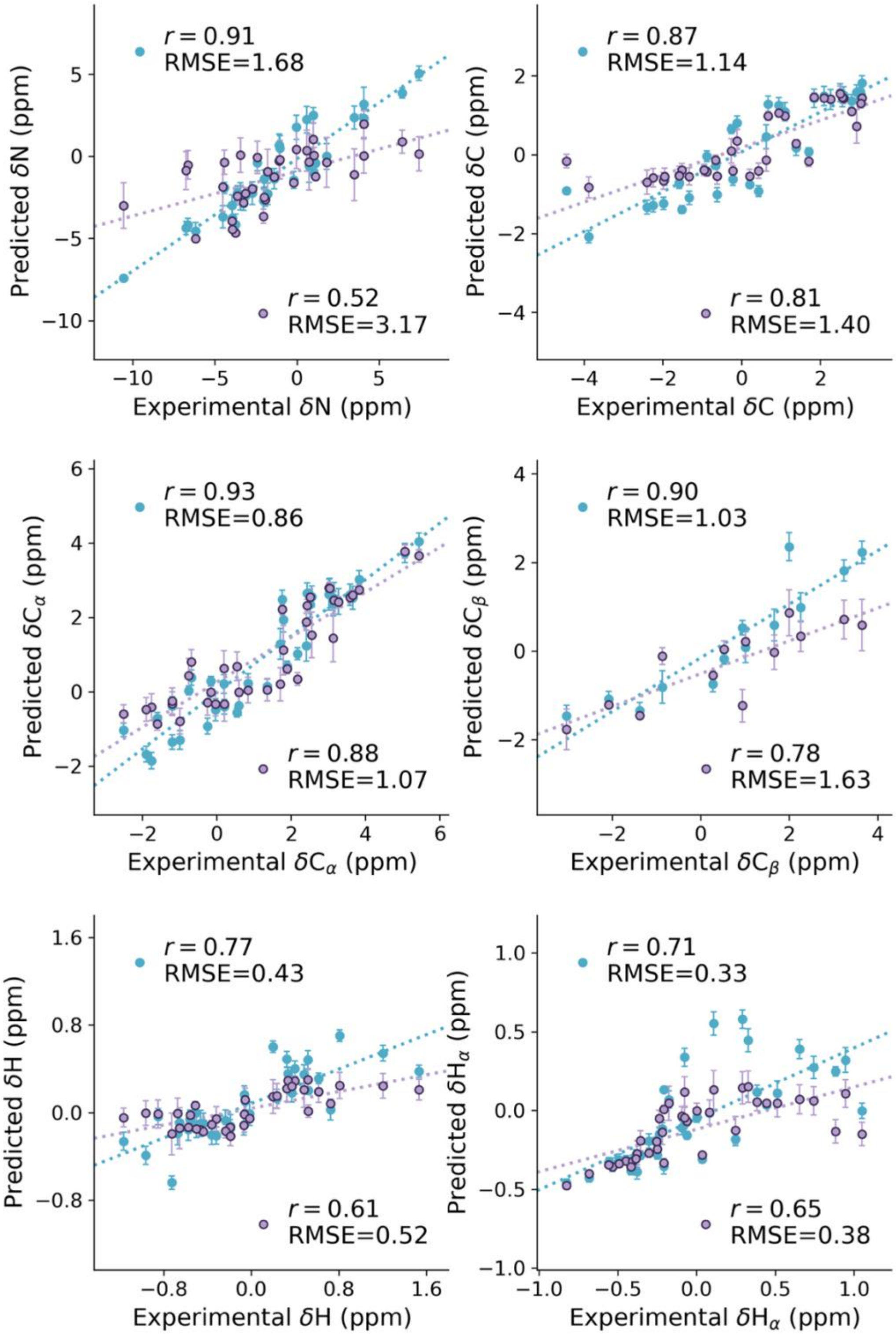


**Figure S1 | Agreement between experimental and predicted chemical shifts.** Correlation between experimental and computed CS for the backbone atoms from simulations performed on a de novo designed mini-protein (PDB ID: 2ND3). In blue simulations performed using Martini3-NMR and in indigo Martini3-DSSP.

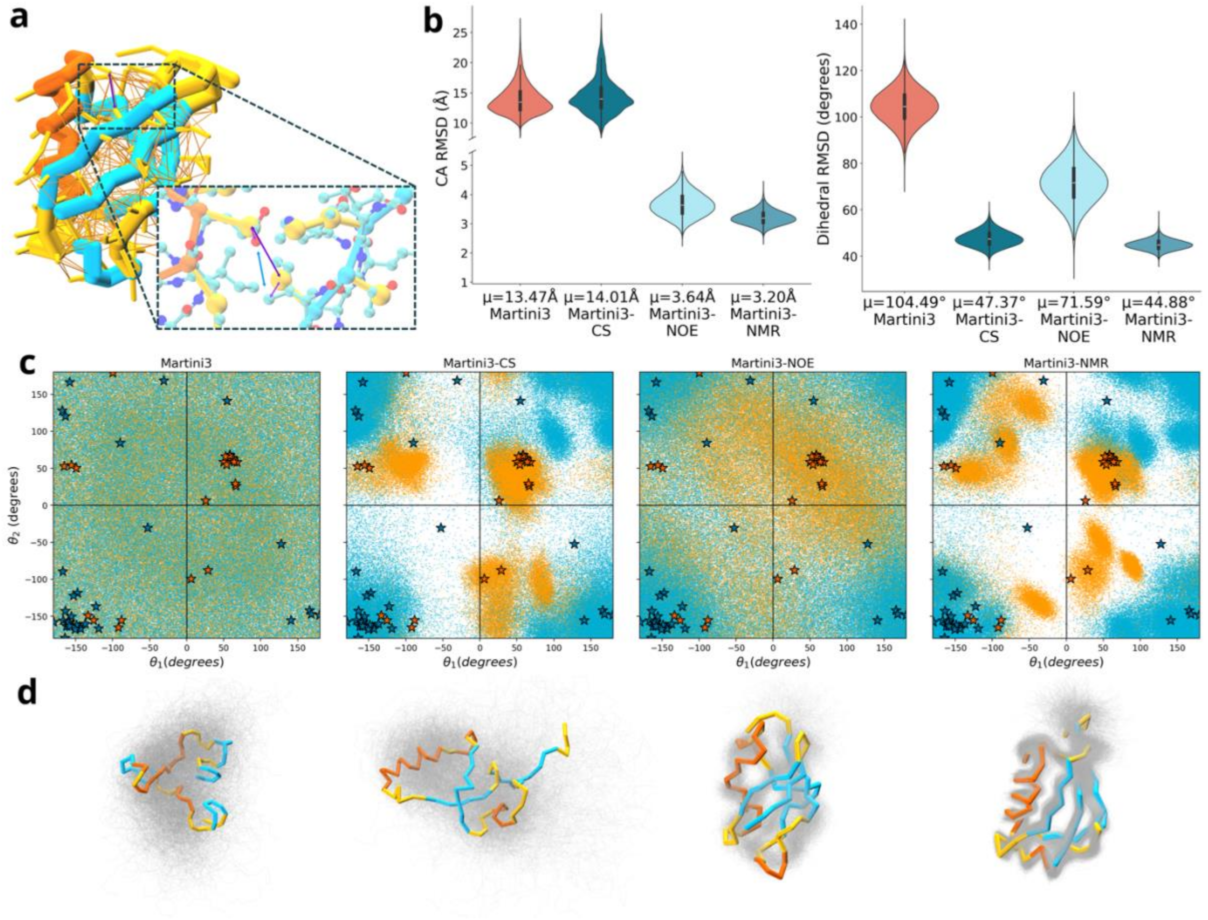


**Figure S2. | Effect of CS and NOE restraints in keeping tertiary packing.** (a) Schematic representation of NOEs mapping in Martini3-NMR. Orange lines show the mapping of NOEs restraints across ubiquitin (PDB CODE: 1UBQ). The inset shows the CG structure overlaid to an all-atom representation, where NOE restraints are represented by arrows between CG beads mapped on the side chain center of mass. (b) Cα (left) and dihedral (right) root mean square deviation (RMSD) distributions (as violin plots) for simulations of ubiquitin carried out using unrestrained Martini3, only CS restraints (Martini3-CS), only NOE restraints (Martini3-NOE) or CS and NOE restraints (Martini3-NMR). (c) From left to right: pseudo-Ramachandran plots in $\theta_1$ and $\theta_2$ dihedral space describing backbone secondary structure in simulations carried out using unrestrained Martini3, only CS restraints (Martini3-CS), only NOE restraints (Martini3-NOE) or CS and NOE restraints (Martini3-NMR). (d) From left to right, Ubiquitin ensembles obtained from simulations carried out using unrestrained Martini3, only CS restraints (Martini3-CS), only NOE restraints (Martini3-NOE) or CS and NOE restraints (Martini3-NMR).

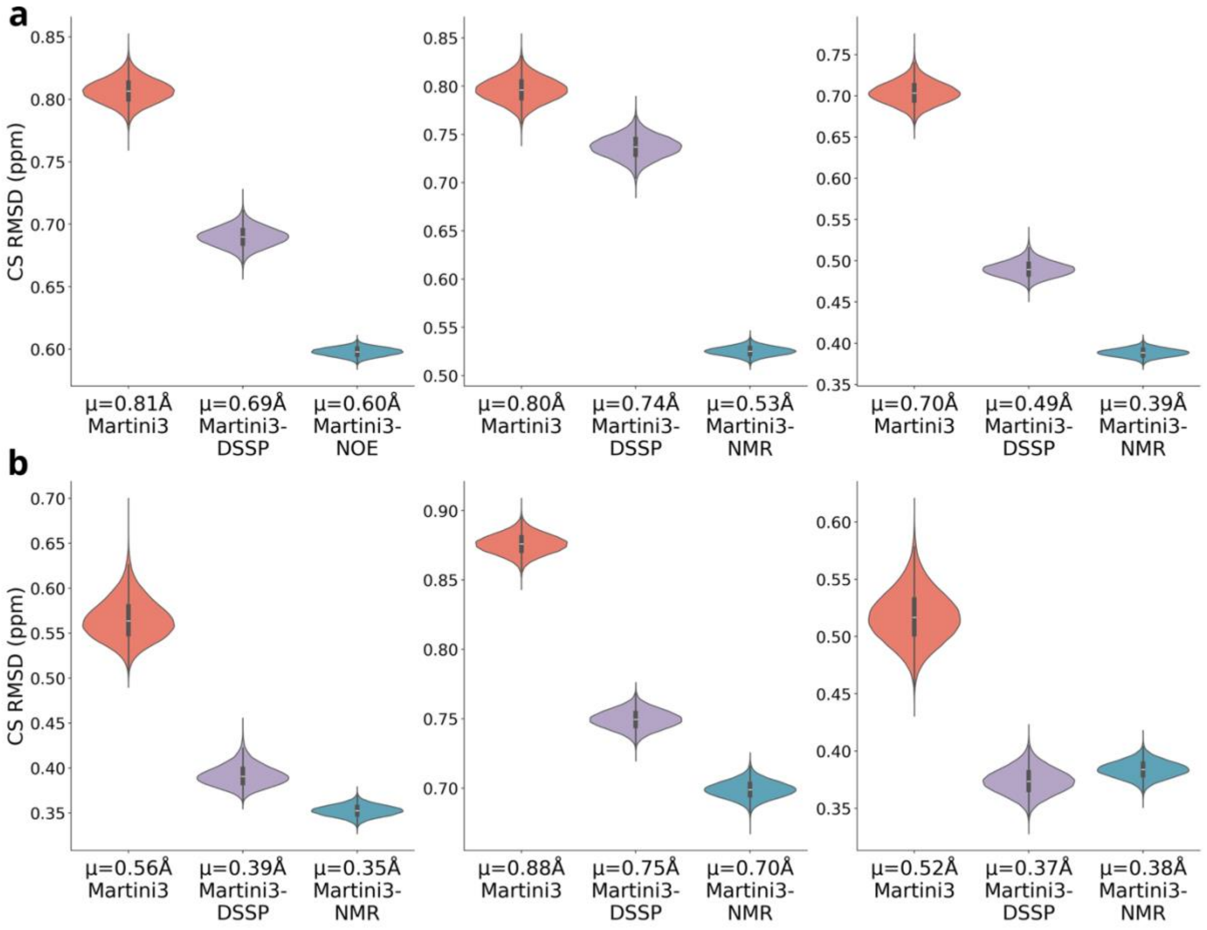


**Figure S3. Chemical shift RMSD obtained from simulations of soluble and membrane proteins.** (a) Chemical shift RMSD of simulations of soluble proteins. From left to right are KRAS (PDB ID: 7KYZ), the SH3 tandem domains of the human KIN protein (PDB ID: 9COJ) and a human leptin (PDB ID: 8K6Z). (b) Chemical shift RMSD of simulations of membrane proteins. From left to right are the single-span transmembrane helical domains of the human tyrosine kinase ErbB1 (PDB ID: 2M0B), the transmembrane anchor domain of the bacterial autotransporter YadA (PDB ID: 2LME), and the phospholamban pentamer (PDB ID: 2KYV). The distributions reflect the simulations performed with Martini3 (unrestrained, pink salmon), Martini3-DSSP (indigo) and Martini3-NMR (teal).

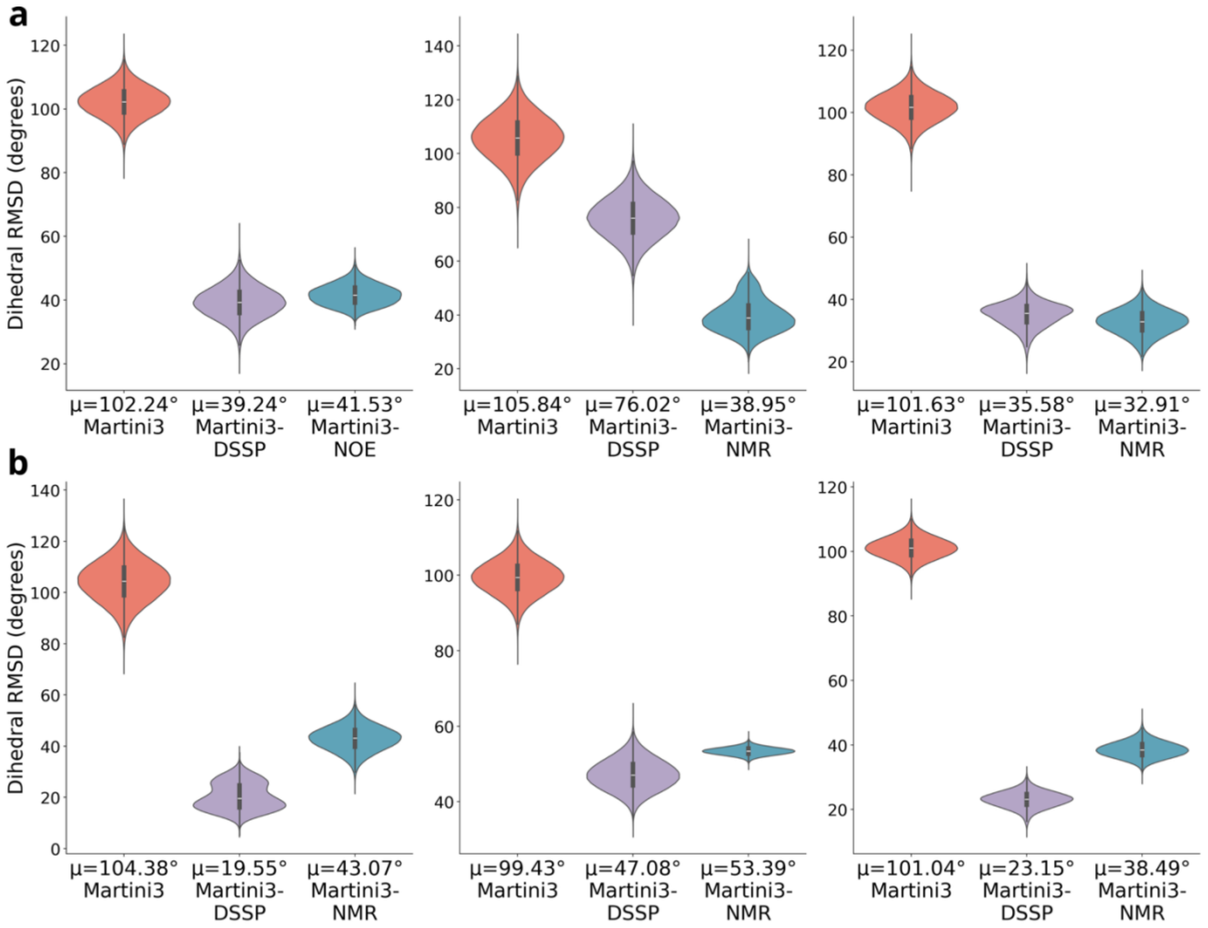


**Figure S4 | Dihedral RMSD obtained from simulations of soluble and membrane proteins.** (a) Dihedral RMSD of simulations of soluble proteins. From left to right are KRAS (PDB ID: 7KYZ), the SH3 tandem domains of the human KIN protein (PDB ID: 9COJ) and a human leptin (PDB ID: 8K6Z). (b) Dihedral RMSD of simulations of membrane proteins. From left to right are the single-span transmembrane helical domains of the human tyrosine kinase ErbB1 (PDB ID: 2M0B), the transmembrane anchor domain of the bacterial autotransporter YadA (PDB ID: 2LME), and the phospholamban pentamer (PDB ID: 2KYV). The distributions reflect the simulations performed with Martini3 (unrestrained, pink salmon) , Martini3-DSSP (indigo) and Martini3-NMR (teal).

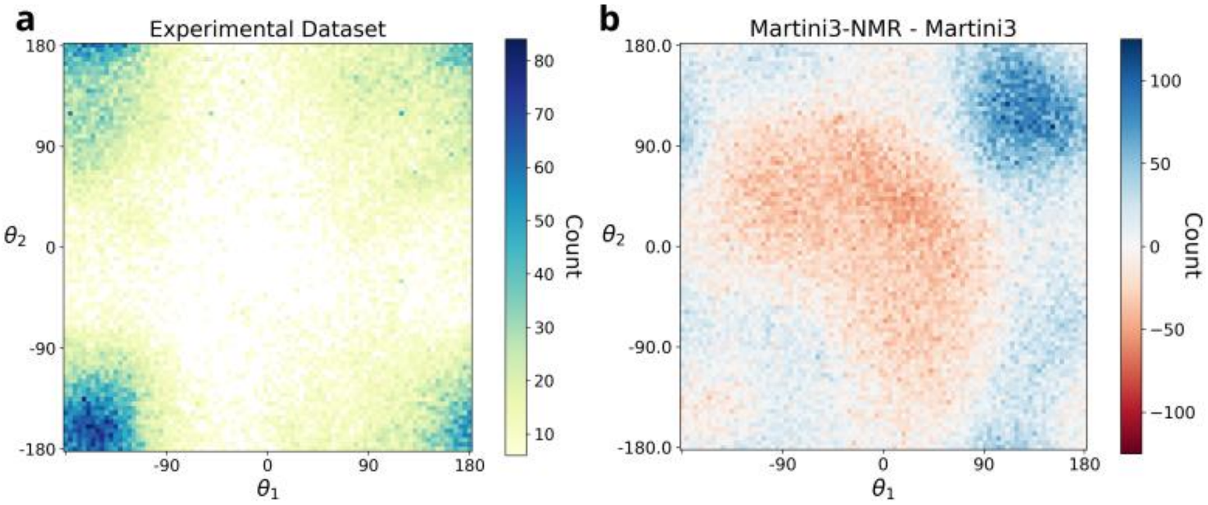


**Figure S5 | Dihedral angles in disordered regions of experimental structures and Martini3-NMR leptin ensembles.** (a) Pseudoramachandran plot composed of $\theta_1$ and $\theta_2$ angles (Figure 1a) and calculated using the structures of loop regions in folded proteins as employed to train Napshift-CG. (b) Difference between the pseudoramachandran plots computed using Martini3-NMR ensemble of Leptin (disordered loop regions) with respect to Martini3 (unrestrained). Blue points indicate regions of the pseudoramachandran sampled more by Martini3-NMR, while red coloring suggests those regions sampled more by Martini3 unrestrained. The latter fall into forbidden regions of the pseudoramachandran plot as observed in experimental structures, indicating that the NMR restraints have significantly improved the dihedral space with respect to unrestrained simulations.

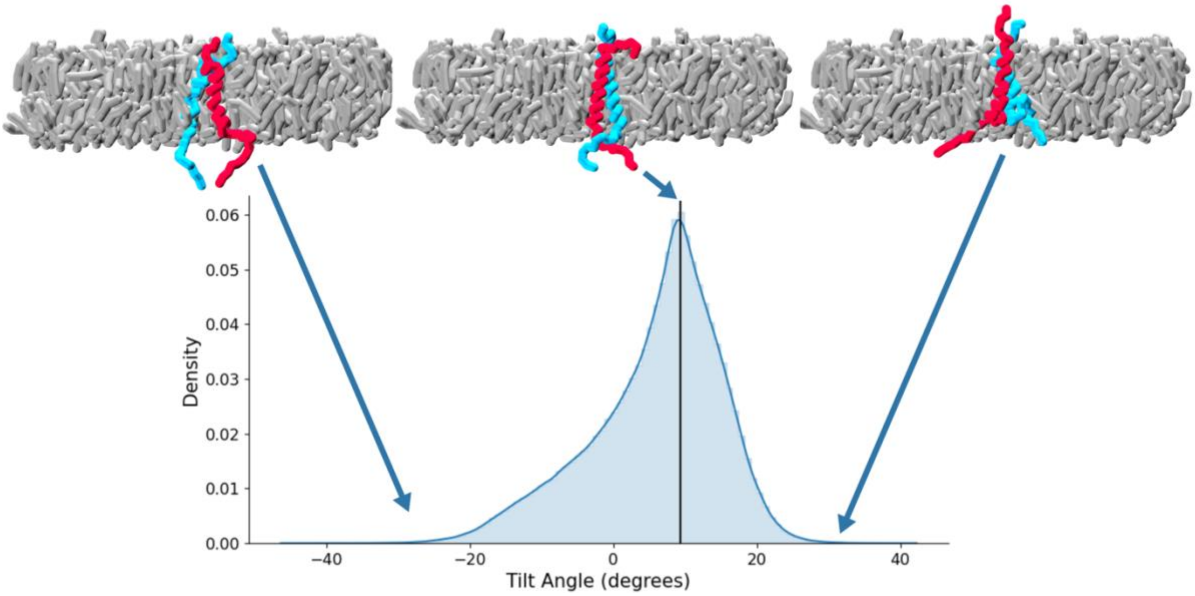


**Figure S6 | Orientation of the transmembrane helices of the human tyrosine kinase ErbB1 as sampled by Martini3-NMR.** Distribution of the tilt angle describing the relative orientation of the TM helices as sampled by Martini3-NMR. In the top panels ensembles reflective of low, median and high tilt angles as pointed by the arrow. Helices are shown in cyan and red while the membrane is shown in gray.

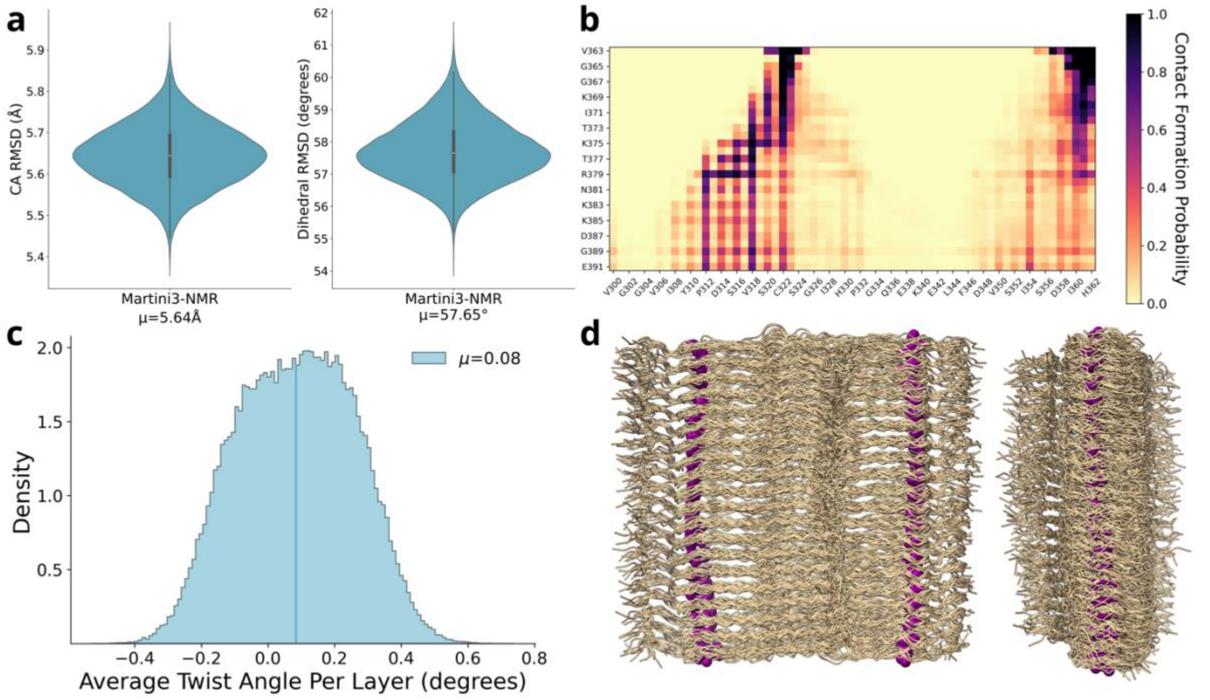


**Figure S7 | NMR restrained simulations of the tau amyloid core show minimal twisting when compared to unrestrained simulations.** (a) $C_\alpha$ (left) and dihedral (right) RMSD distributions (as violin plots) of the simulations of the tau amyloid (PDB ID: 8G58) carried out either using Martini3-NMR. (b) Fraction of contacts between the intrinsically disordered regions and the core. (c) Distributions of the twisting angle per layer from simulations performed using Martini3-NMR (steel blue). (d) Ensemble of the amyloid core obtained from simulations performed using Martini3-NMR, with the BB beads at the N- and C-terminal residues highlighted as magenta spheres to better track twisting.
